# Inferring microbial co-occurrence networks from amplicon data: a systematic evaluation

**DOI:** 10.1101/2020.09.23.309781

**Authors:** Dileep Kishore, Gabriel Birzu, Zhenjun Hu, Charles DeLisi, Kirill S. Korolev, Daniel Segrè

## Abstract

Microbes tend to organize into communities consisting of hundreds of species involved in complex interactions with each other. 16S ribosomal RNA (16S rRNA) amplicon profiling provides snapshots that reveal the phylogenies and abundance profiles of these microbial communities. These snapshots, when collected from multiple samples, have the potential to reveal which microbes co-occur, providing a glimpse into the network of associations in these communities. The inference of networks from 16S data is prone to statistical artifacts. There are many tools for performing each step of the 16S analysis workflow, but the extent to which these steps affect the final network is still unclear. In this study, we perform a meticulous analysis of each step of a pipeline that can convert 16S sequencing data into a network of microbial associations. Through this process, we map how different choices of algorithms and parameters affect the co-occurrence network and estimate steps that contribute most significantly to the variance. We further determine the tools and parameters that generate the most accurate and robust co-occurrence networks based on comparison with mock and synthetic datasets. Ultimately, we develop a standardized pipeline (available at https://github.com/segrelab/MiCoNE) that follows these default tools and parameters, but that can also help explore the outcome of any other combination of choices. We envisage that this pipeline could be used for integrating multiple data-sets, and for generating comparative analyses and consensus networks that can help understand and control microbial community assembly in different biomes.

**Importance:** To understand and control the mechanisms that determine the structure and function of microbial communities, it is important to map the interrelationships between its constituent microbial species. The surge in the high-throughput sequencing of microbial communities has led to the creation of thousands of datasets containing information about microbial abundances. These abundances can be transformed into networks of co-occurrences across multiple samples, providing a glimpse into the structure of microbiomes. However, processing these datasets to obtain co-occurrence information relies on several complex steps, each of which involves multiple choices of tools and corresponding parameters. These multiple options pose questions about the accuracy and uniqueness of the inferred networks. In this study, we address this workflow and provide a systematic analysis of how these choices of tools and parameters affect the final network, and on how to select those that are most appropriate for a particular dataset.

## Introduction

Microbial communities are ubiquitous and play an important role in marine and terrestrial environments, urban ecosystems, metabolic engineering, and human health [1, 2]. These microbial communities, or microbiomes, often comprise several hundreds of different microbial strains interacting with each other and their environment, often through intricate metabolic and signaling relationships. Understanding how these interconnections shape community structure and functionalities is a fundamental challenge in microbial ecology, with applications in the study of microbial ecosystems across different biomes. With the advancement in DNA sequencing technologies [3] and data processing methods, more information can be extracted from these microbial community samples than ever before. In particular, high-throughput sequencing, including community metagenomic sequencing and sequencing of 16S rRNA gene amplicons, has the potential to help detect, identify and quantify a large portion of the constitutive microorganisms of a microbiome [4, 5]. These advances have led to large-scale data collection efforts involving environmental (Earth Microbiome Project) [2], marine (Tara Oceans Project) [6] and human-associated microbiota (Human Microbiome Project) [7].

This wealth of information on the composition and functions of a community at different times and under different environmental conditions has the potential to help us understand how communities assemble and operate. A powerful tool for translating microbiome data into knowledge is the construction of possible inter-dependence networks across species. The importance of these networks of relationships is two fold: first, such networks can serve as maps that help identify hubs of keystone species [8, 9], or basic microbiome changes that occur as a consequence of environmental perturbations or underlying host conditions [10]; second, networks of inter-dependencies can serve as a key bridge towards building mechanistic models of microbial communities, greatly enhancing our capacity to understand and control them. For example, multiple studies have shown the importance of specific microbial interactions in the healthy microbiome [5] and others have shown how changes in these interactions can lead to dysbiosis [11, 10, 12]. In the context of terrestrial bio-geochemistry, co-occurrence networks have been proposed as a valuable approach towards reconstructing the processes leading to microbiome assembly [13], and understanding the response of microbial communities to environmental perturbations [14].

Direct high-throughput measurement of interactions, e.g. through co-culture micro-droplet experiments [15, 16], or spatial visualization of natural communities [17] is possible, but it requires specific technological capabilities, and has yet to be extensively used. In parallel, sequencing data across multiple samples can be used for estimating co-occurrence relationships between taxa. While the the relationship between directly measured interactions and statistically inferred co-occurrence is still poorly understood [18], a significant amount of effort has gone into estimating correlations from large microbiome sequence datasets. Co-occurrence networks have microbial taxa as nodes, and edges that represent the frequent co-occurrence (or negative correlations) across different datasets.

One of the most frequently used avenues for inferring co-occurrence networks is the parsing and analysis of 16S sequencing data [9, 19]. A large number of software tools and pipelines have been developed to analyze 16S sequencing data, often focused on addressing the many known limitations of this methodology, including resolution, sequencing depth, compositional nature, sequencing errors and copy number variations [20, 21]. Popular methods for different phases of the analysis of 16S data include tools for: (i) denoising and clustering sequencing reads [22, 23]; (ii) assigning taxonomy to the reads [24, 25]; (iii) processing and transforming the taxonomy count matrices [26]; and (iv) inferring the co-occurrence network [27, 28]. Different specific algorithms are often aggregated into popular platforms (like MG-RAST [29], Qiita [30]) and packages (such as QIIME [22]) that provide pipelines for 16S data analysis. The different methods and tools developed to solve issues arising in 16S analysis can lead to vastly different inferences of community compositions and co-occurrence networks [31, 32], making it difficult to reliably compare networks across different publications and studies. This is partially due to the fact that existing platforms are typically focused on Operational Taxanomic Unit (OTU) generation and not on the effects of upstream statistical methods on the inferred co-occurrence networks. Furthermore, no organized framework currently exist to systematically analyze and compare existing components of the data analysis from amplicons to networks. More broadly, given the lack of comprehensive comparisons between directly observed microbial interactions (e.g. from co-culture experiments) and co-occurrence networks, there is no straightforward way to determine which set of tools or methods generate the most accurate networks.

In this study, we present a standardized 16S data analysis pipeline called Microbial Co-occurrence Network Explorer (MiCoNE) that produces robust and reproducible co-occurrence networks from community 16S sequence data, and allow users to interactively explore how the network would change upon using different alternative tools and parameters at each step. Our pipeline is coupled to an online integrative tool for the organization, visualization and analysis of inter-microbial networks. In addition to making this tool freely available, we implemented a systematic comparative analysis to determine which steps of the pipeline have the largest influence on the final network, and what choice seems to provide best agreement with the tested mock and synthetic datasets. We believe that these steps will ensure better reproducibility and easier comparison of co-occurrence networks across datasets. We expect that our tool will also be useful for benchmarking future alternative methods, and for ensuring a transparent evaluation of the possible biases introduced by the use of specific tools.

## Results

### Microbial Co-occurrence Network Explorer (MiCoNE)

We have developed MiCoNE, a flexible and modular pipeline for 16S amplicon sequencing rRNA data (hereafter mentioned simply as 16S data) analysis, that allows us to infer microbial co-occurrence networks. It incorporates various popular, publicly available tools as well as custom Python modules and scripts to facilitate inference of co-occurrence networks from 16S data (see Methods). Using MiCoNE one can obtain co-occurrence networks by applying to 16S data (or to already processed taxonomic count matrices) any combination of the available tools. The effects of changing any of the intermediate step can be monitored and evaluated in terms of its final network outcome, as well as on any of the intermediate metrics and data outputs. The MiCoNE pipeline workflow is shown in Figure 1. The different steps for going from 16S data to co-occurrence networks can be grouped into four major modules; (i) the denoising and clustering (DC) step, which handles denoising of the raw 16S sequencing data into representative sequences; (ii) the taxonomy assignment (TA) step that assigns taxonomic labels to the representative sequences; (iii) the OTU processing (OP) step that filters and transforms the taxonomy abundance table; and finally (iv) the network inferences (NI) step which infers the microbial co-occurrence network. Each process in the pipeline supports alternate tools for performing the same task (see Methods and Figure 1). A centralized configuration file contains all the specifications for what modules are used in the pipeline, and can be modified by the user to choose the desired set of tools. In what follows, we perform a systematic analysis of each step of the pipeline to estimate how much the final co-occurrence network depends on the possible choices at each step. We also evaluate a large number of tool combinations to determine a set of recommended default options for the pipeline and provide the users with a set of guidelines to facilitate tool selection as appropriate for their data.

**Figure 1:**
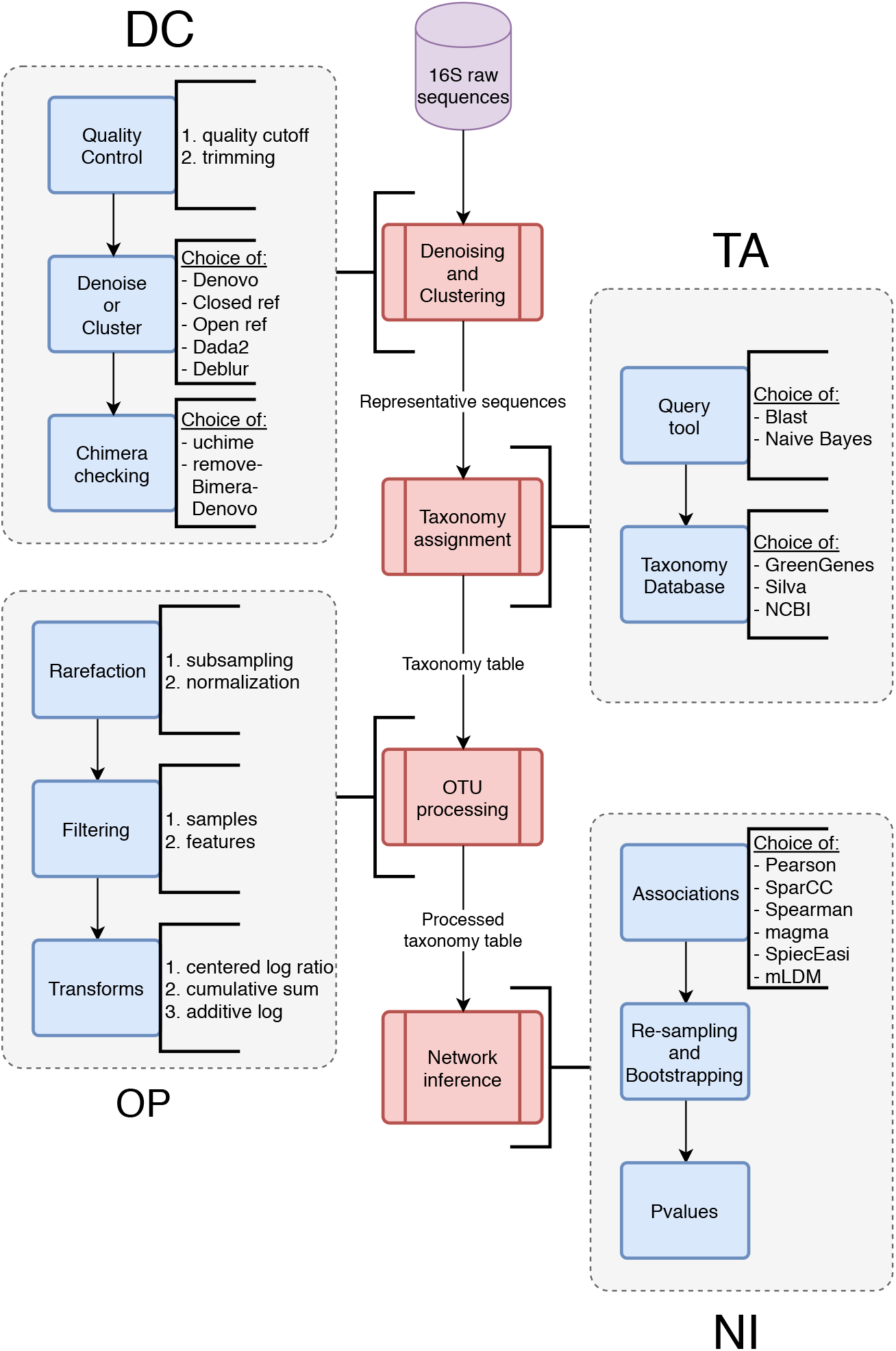
The workflow of the microbial co-occurrence analysis pipeline. The steps can be grouped into four major groups: **(DC) D**enoising and **C**lustering, **(TA) T**axonomy **A**ssignment, **(OP) O**TU or ESV **P**rocessing, and **(NI) N**etwork **I**nference. Each step incorporates several processes, each of which in turn have several alternate algorithms for the same task (indicated by the text to the right of the blue boxes). The text along the arrows describes the data that is being passed from one step to another. For details on each process and data types, see Methods.

Our analysis involves two types of data: The first type consists of sets of 16S sequencing data from real communities sampled from human Stool and Oral microbiomes. The second are datasets synthetically or artificially created for the specific goal of helping evaluate computational analysis tools (see Methods). In particular, in order to objectively compare, to the extent possible, how well each step in MiCoNE best captures the underlying data, we use both mock data (labelled mock4, mock12 and mock16) from mockrobiota [33] as well as, synthetically generated reads from an Illumina read simulator called ART [34]. These mock datasets consist of fake sequencing reads generated from reads obtained from synthetic microbial isolates mixed in know proportions. They contain the expected compositions along with the reference sequences for the organisms in the mock community. The synthetic reads were simulated using three different taxonomy distribution profiles, namely soil and water microbiomes obtained Earth Microbiome Project (EMP) [2] and Stool microbiome that is used in our real community analysis [35]. Reference sequences were generated using National Center for Biotechnology Information (NCBI) and the Decard package [31] for these taxonomy profiles. Detailed information on the mock communities and the settings used to generate the synthetic data are provided in the Methods section.

### The choice of reference database has the biggest impact on inferred networks

In order to analyze the effect of different statistical methods on the inferred co-occurrence networks, we generated co-occurrence networks using all possible combinations of methods and estimated the variability in the networks due to each choice (Figure 1). This analysis is performed while keeping the network inference algorithm (NI step) the same throughout the analysis. The effects of various steps on the final co-occurrence network is estimated by building a linear model of the edges of the network as a function the various step in the analysis pipeline (see Methods). Figure 2B, shows the fraction of total variation among the co-occurrence networks due to the first three steps of the pipeline. In other words, each point corresponds to a different combination of tools, and captures how much the final network is affected by such choice. The 16S reference database contributes the most (∼ 25%) to variation in the networks. This is also reflected in the fact that the networks can be clearly separated based on the database used (Figure 2B). This indicates that the taxonomy assigned to the reference sequences drastically alters the co-occurrence network. In fact the variability induced by taxonomy assignment is much more significant than that due to the variability induced based on how the reference sequences themselves are identified (in the DC step). The grouping of the networks by taxonomy assignment into clusters (Figure 2B) seems to derive from the mislabelling of constitutive taxa that are present in high abundance in the community, which drastically alter the nodes and hence the underlying network topology. The residual variation (Figure 2A) can be seen as an artifact that arises when multiple steps are changed at the same time. Another interesting observation (elaborated in detail in the denoising and clustering section) is that the dissimilarity between the networks decreases when the low abundance OTUs are removed from the network. These results suggest that the most important criterion for accurate comparative analyses of co-occurrence networks is the taxonomy reference database.

**Figure 2:**
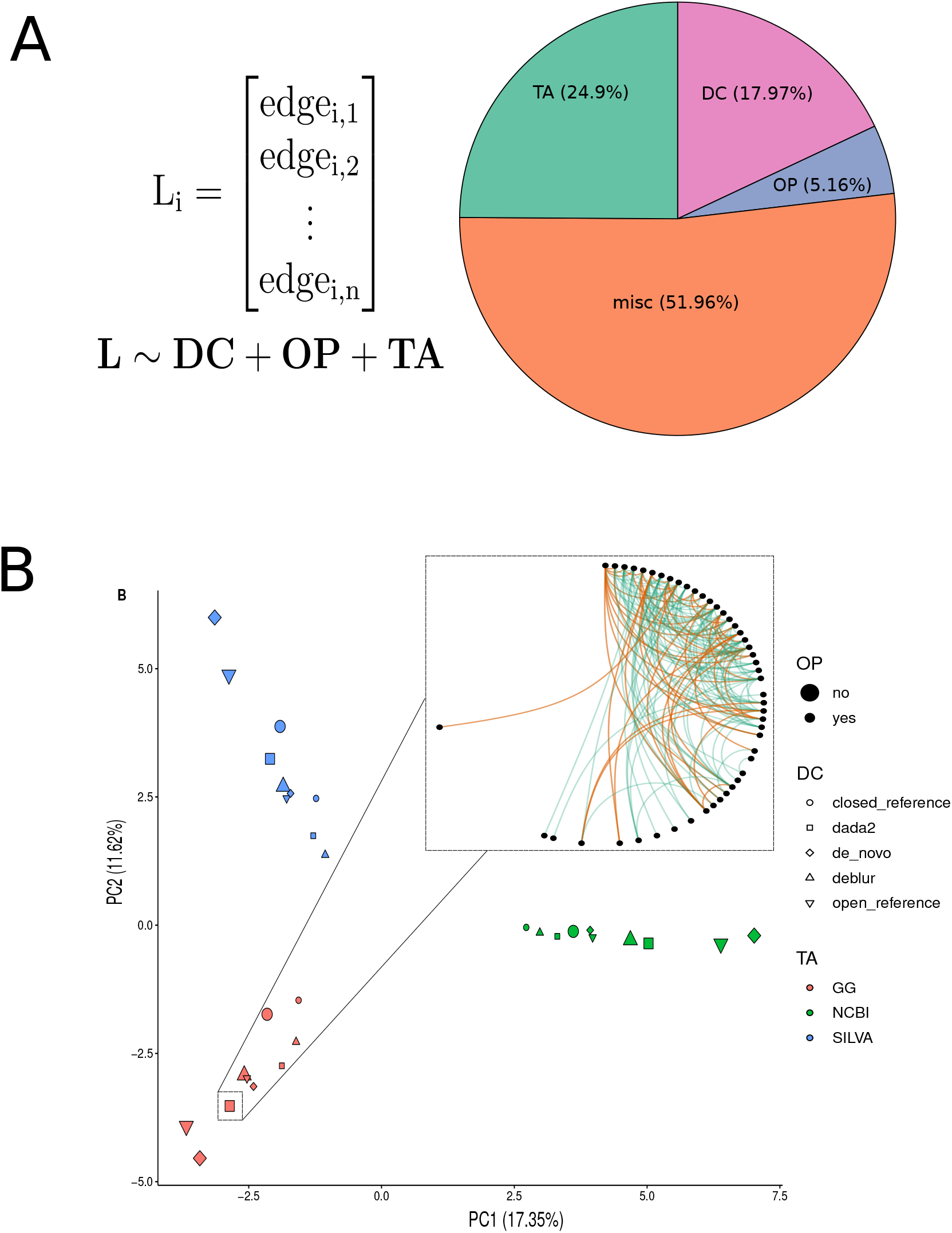
The choice of database contributes to the most variance in the networks. **(A)** The total relative variance in the networks contributed by the DC, TA and OP steps of the pipeline (right) and the linear model used to calculate the relative variance (left), see the Methods section for details. **(B)** All combinations of inferred networks are shown as points on a PCA plot. The color of the points corresponds to the taxonomy database, the shape corresponds to the denoising/clustering method and the size corresponds to whether low abundance OTUs were removed or not. **(B inset)** The network generated using DC=dada2, TA=GG, OP=no and NI=SPARCC and represents the particular point shown (big red square). The plot clearly shows that the points can be separated based on the TA step and that the differences due to the DC and OP steps are not as significant.

### Denoising and clustering methods differ in their identification of less common reference sequences

Denoising and clustering are commonly carried out to generate representative sequences from the raw 16S sequencing data and to obtain the OTU/Exact Sequence Variant (ESV) tables (counts of these representative sequences for each sample). In order to compare the OTU tables generated by different tools we processed the same 16S sequencing reads (healthy samples from a fecal microbiome transplant study [35]) using 5 different methods: open-reference clustering, closed-reference clustering, denovo clustering, Divisive Amplicon Denoising Algorithm 2 (DADA2) [23] and Deblur [36]. The first three methods are from the Quantitative Insights Into Microbial Ecology 1 (QIIME1) [22] package. We find that there is good agreement in the OTU/ESV tables when different combinations of methods are used to generate them (Supplementary Figure S1).

To compare the representative sequences generated by these methods we employ both the weighted [37] (Figure 3A) and unweighted UniFrac method [38] (Figure 3B). The weighted UniFrac distance metric takes into account the counts of the representative sequences, whereas the unweighted UniFrac distance metric does not and hence gives equal weights to each sequence. From Figure 3A one can see that the representative sequences generated by the different methods are similar to each other when weighted by their abundance. Figure 3B on the other hand shows an increase in dissimilarity between each pair of methods suggesting that the methods might differ in the treatment of sequences of low abundance. In order to verify this claim, for each of these methods we use the Greengenes (GG) taxonomy database to assign taxonomies to the representative sequences. We then correlate the abundances of matching taxonomies between a pair of DC methods (Figure S1A and B). The ESV tables generated by methods that perform denoising are very similar to each other (∼ 0.91) and the OTU tables generated by the clustering methods are very similar to each other (∼ 0.9), but results of denoising and clustering are highly uncorrelated with each other (∼ 0.4) (Figure S1C).

**Figure 3:**
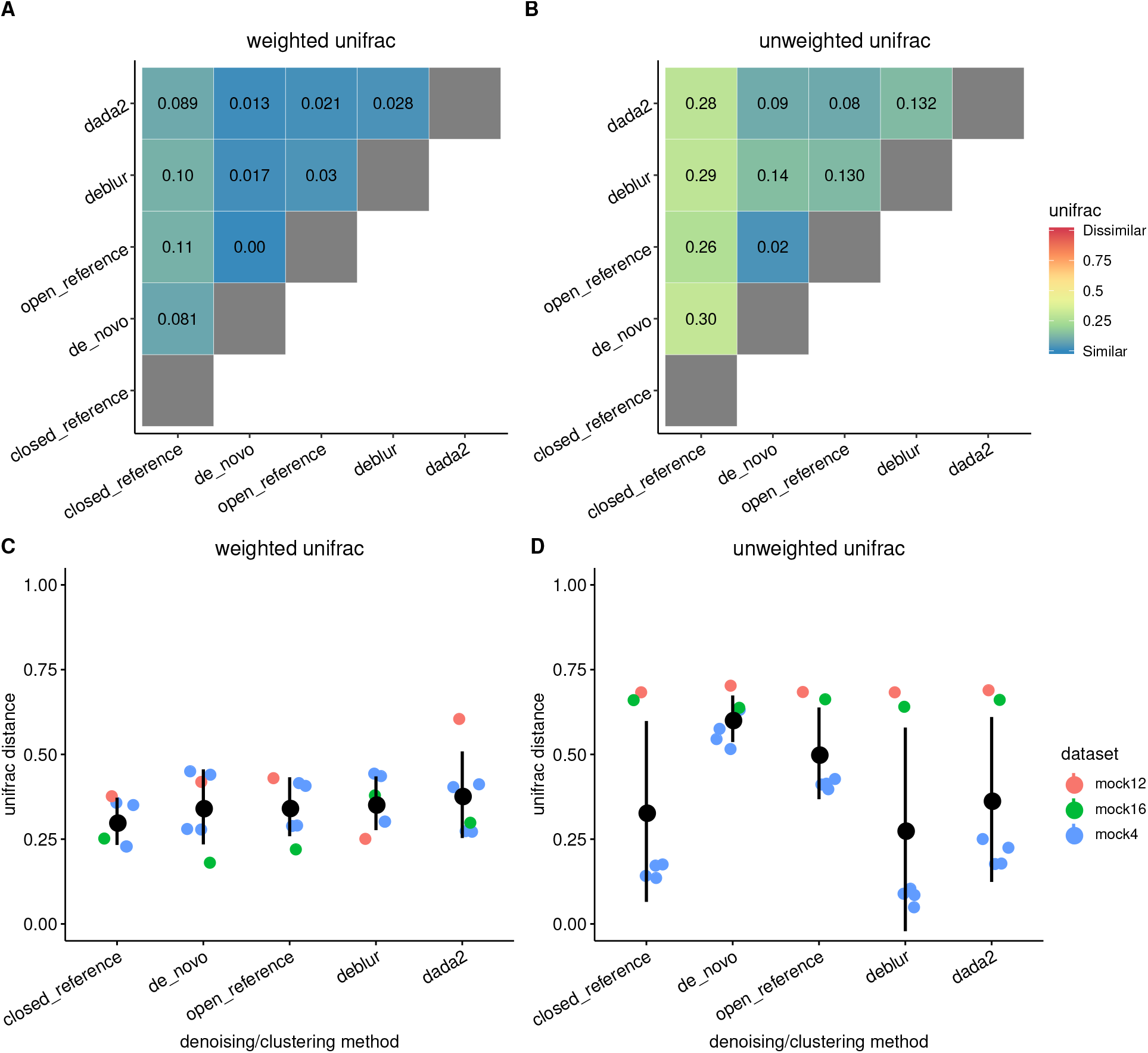
The representative sequences generated by the different denoising/clustering methods are very similar but differ in the sequences that are in low abundance. **(A)** The average weighted UniFrac distance between the representative sequences shows that the representative sequences and their compositions are fairly identical between the methods, **(B)** The relatively larger average unweighted UniFrac distance indicates that methods differ in their identification of sequences of low abundance, **(C, D)** The distributions of the average weighted UniFrac distance between the expected sequence profile and the calculated sequence profile in mock datasets. **(D)** The distributions of the average unweighted UniFrac distance show that dada2 and Deblur were the best performing methods in most of the datasets.

These comparisons only elucidate the pairwise similarity or dissimilarity of a pair of methods. In order to determine the tool that most accurately recapitulates the reference sequences in the samples, we used the 16S sequences from the mock datasets. In particular, we used the pipeline to process mock community datasets using each of the possible methods included for this step. We next compared predicted representative sequences with expected representative sequences and their distribution. The results (Figure 3C and D) show that, for the mock datasets, the different methods perform similar to each other, exactly as observed in the case of the real dataset. However, the mock predicted sequence distributions are substantially different from the expected sequence distribution. This result is more exaggerated in the case of the unweighted UniFrac metric, where some of the datasets show a very high deviation from the expected sequences. These high deviations are primarily in two of the three datasets that were analyzed and show that the datasets themselves play a big role in the performance of these methods. This can be clearly seen in the performance (weighted UniFrac distance) of DADA2 and Deblur on mock12 and mock16 datasets, where, Deblur outperforms DADA2 on mock12 but the under-performs on mock16.

There is no method that clearly outperforms the rest in all datasets. Based on their slightly better performance on the mock datasets, their *de novo* error correcting nature and other previous studies [39], DADA2 and Deblur seem to be in general the most reliable. Given the unexpected poor performance of Deblur on the synthetic data, the default algorithm in the pipeline was chosen to be DADA2 (Supplementary Figure S3).

### Taxonomy databases vary widely in taxonomy hierarchy and update frequency

Taxonomy databases are used to assign taxonomic identities to the representative sequences obtained after the DC step. In order to compare the assigned taxonomies from different databases, we use the same reference sequences and assign taxonomies to them using different taxonomy reference databases. The three 16S taxonomic reference databases used in this study are SILVA [25], GG [24] and NCBI RefSeq [40]. SILVA and GG are two popular 16S databases used for taxonomy identification. The NCBI RefSeq nucleotide database contains 16S rRNA sequences as a part of two BioProjects - 33175 and 33317. The three databases vastly differ in terms of their last update status - GG was last updated on May 2013, SILVA was last updated on December 2017 at the time of writing and NCBI is updated as new sequences are curated. Since updates to taxonomic classifications are frequent, these databases vary significantly in terms of taxonomy hierarchies including species names and phylogenetic relationships [41].

The representative sequences obtained from the DADA2 method in DC step were used for taxonomic assignment using the three reference databases. Figure 4A depicts a flow diagram that shows how the top 50 representative sequences (sorted by abundance) are assigned a Genus according to the three different databases. We observe that not only does the assigned Genus composition vary significantly, but the percentage of unassigned representative sequences (gray) also differ. Even the most abundant representative sequence is assigned to an “unknown” Genus in two of the three databases. A representative sequence might be assigned an “unknown” Genus for one of two reasons: the first is if the taxonomy identifier associated with the sequence in the database did not contain a Genus; the second (more likely) reason is that the database contains multiple sequences that are very similar to the query (representative) sequence and the consensus algorithm (from Quantitative Insights Into Microbial Ecology 2 (QIIME2)) is unable to assign one particular Genus at the required confidence. After assigning all the representative sequences to taxonomies we perform a pairwise comparison of the similarity between assignments from different databases at every taxonomic level (Figure 4B). The assignments beyond Family level (Family, Genus and Species) are very dissimilar with < 70% similarity between any pair of databases. There are no two reference databases that are more similar than the other pairs, with GG and SILVA producing only marginally similar assignments compared to NCBI. This implies that the taxonomy assignments from each reference database are fairly unique and are largely responsible for the differences observed in the co-occurrence networks generated from different taxonomy databases.

Supplementary Figure S4 shows that the top 20 most abundant genera in the three resulting taxonomy composition tables are different. For example, the most abundant genus in the GG taxonomy table was *Escherichia* whereas in the SILVA taxonomy table it was *Escherichia-Shigella*. Although these are minor differences, when comparing a large number of taxonomy composition tables these problems are hard to diagnose.

**Figure 4:**
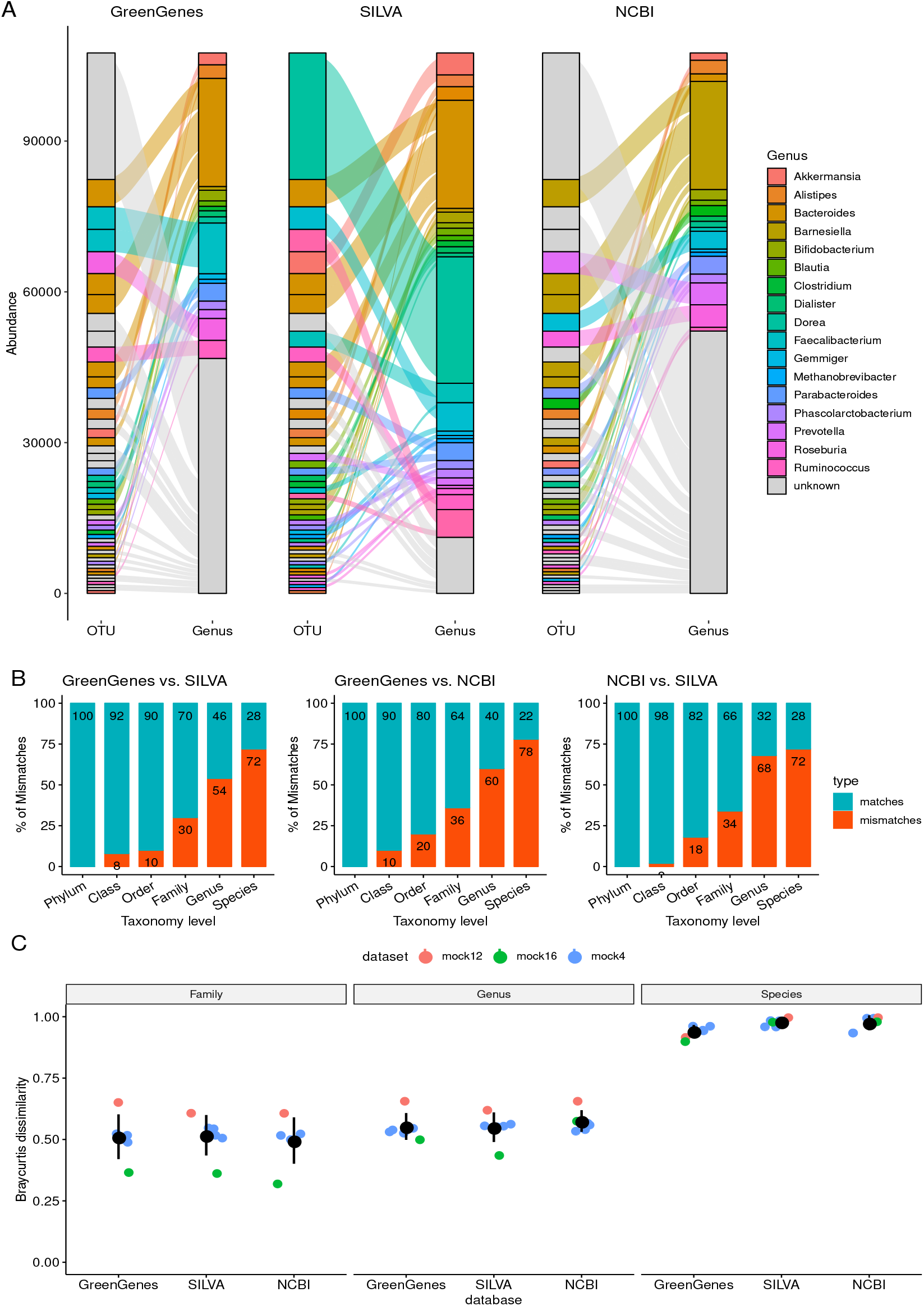
Taxonomic reference databases vary widely in terms of their taxonomy assignments. **(A)** The assignment of the top 50 representative sequences to their respective taxonomies using the three different reference databases shows how the same sequences are assigned to different Genus. **(B)** The percentage of OTUs assigned to the same taxonomic label when using different reference databases. The percentage of mismatches decrease at higher taxonomic levels but even at the Phylum level there exists around 10% of mismatches. **(C)** The Bray-Curtis dissimilarity between the expected taxonomy profile and calculated taxonomy profile in the mock datasets shows that there is no singular best choice of database for every dataset.

As in the previous section, these comparisons only indicate similarity or dissimilarity between methods. In order to obtain an absolute measure of accuracy of the taxonomic assignments we use the expected reference sequences from the mock datasets as the query sequences for the databases and the expected taxonomic composition as the standard to compare against (Figure 4C). Again, we observe that none of the databases perform better than the others in absolute terms.

Given that no database performs better than others against mock datasets, and that databases are almost equally distant from each other in terms of final output, the choice of which database to use should be driven by other reason. One user-specific way to choose, would be based on the known representation of taxa for the microbiome of interest (see also Discussion). Another reason could be the frequency of updates and the potential for future growth, which prompted us to set NCBI as the MiCoNE standard for taxonomy assignment. In addition to being regularly maintained and updated the NCBI database already has the advantage that its accuracy of assignments is still comparable to the SILVA and GG reference databases that are routinely used as reference databases.

### Networks generated using different network inference methods show notable difference in edge-density and connectivity

The six different network inference methods used in this study are Microbial Association Graphical Model Analysis (MAGMA) [27], metagenomic Lognormal-Dirichlet-Multinomial (mLDM) [42], Sparse InversE Covariance estimation for Ecological Association and Statistical Inference (SpiecEasi) [28], Sparse Correlations for Compositional data (SparCC) [19], Spearman and Pearson. These network inference methods fall into two groups, the first set of methods (Pearson, Spearman, SparCC) infer pairwise correlations while the second set infer direct associations (SpiecEasi, mLDM, MAGMA). Pairwise correlation methods involve calculating the correlation coefficient between every pair of OTU/ESVs leading to the detection of spurious indirect connections. On the other hand, direct association methods use conditional independence to avoid the detection of correlated but indirectly connected OTUs [28, 8].

For the analysis presented in this section, we used the taxonomy composition table obtained using the NCBI reference database as the input for algorithms that infer co-occurrence associations between the microbes. Figure 5A shows the networks inferred from this dataset using the different inference algorithms. The different networks differ vastly in their edge-density and connectivity; even some of the edges in common to these networks have their signs inverted. Note, however, that some of these comparisons depend on the threshold that has to be applied to the pairwise correlations methods (currently 0.3, based on [19]). To get a more quantitative picture of the differences between the inferred networks, we checked the distribution of common nodes and edges (Figure 5B) using UpSet plots [43] (only MAGMA, mLDM, SpiecEasi, SparCC are used in the comparison since Pearson and Spearman add a large number of spurious edges since they are not intended for compositional datasets). The results for the node intersections show that the networks have a large number of nodes in common (63 out of 67 nodes in the smallest network - MAGMA) and no network possesses any unique node. The edge intersections in contrast show that only 19 edges (out of 98 edges in the smallest network - MAGMA) are in common between all the methods and each network has a large number of unique edges. These results indicate that there is a substantial rewiring of connections in the inferred networks.

**Figure 5:**
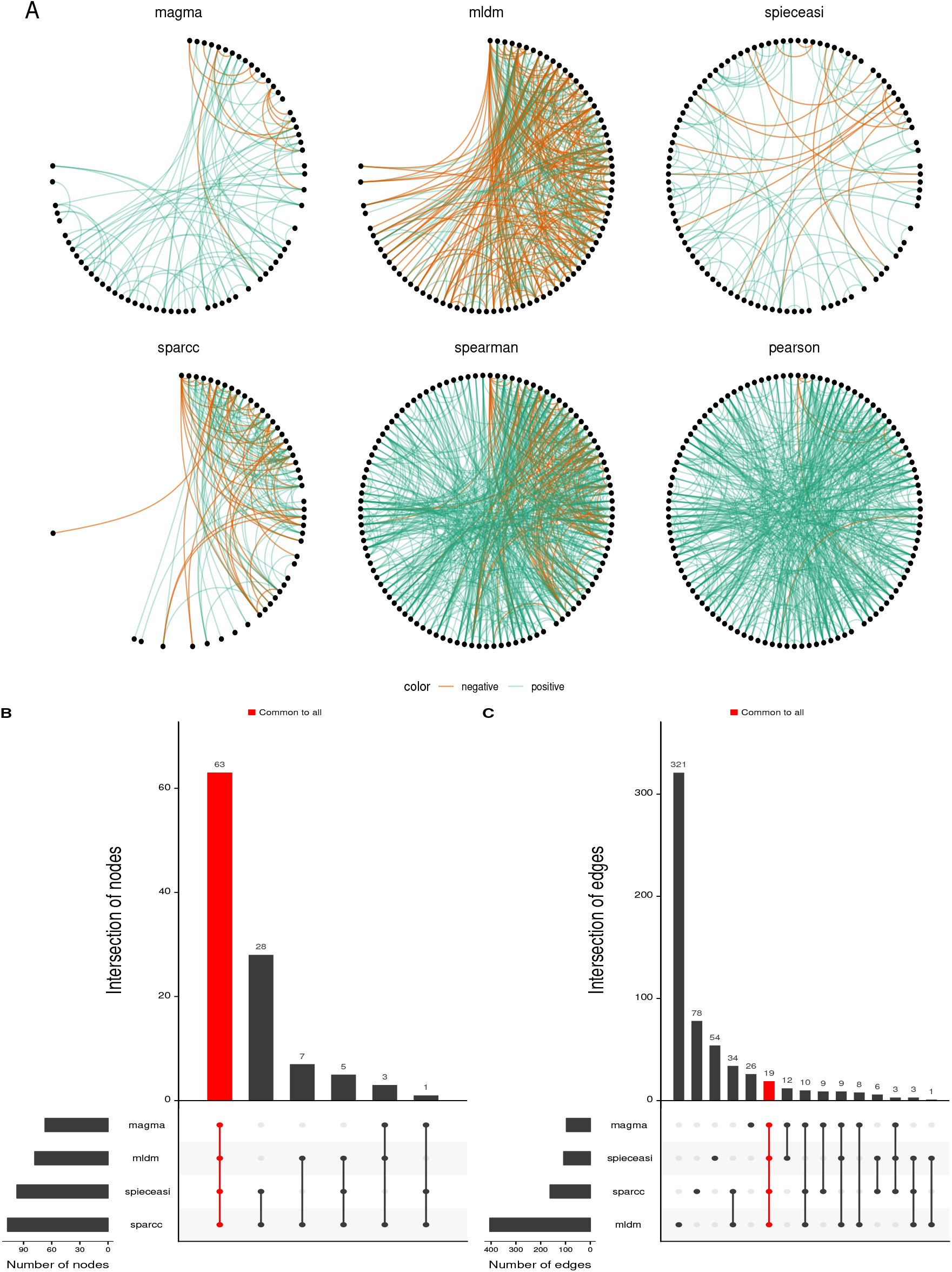
Networks generated using different network inference methods show notable differences both in terms of edge-density and connectivity. **(A)** The six different networks generated by the different network inference methods are very dissimilar. The green links are positive associations and the orange links are negative associations. A threshold of 0.3 was set for the methods that infer pairwise correlations (SparCC, Spearman, Pearson) and no threshold was set for the other methods. **(B)** The node overlap Upset plot [43] indicates that all the networks have a large number of common nodes involved in connections. Whereas, **(C)** The edge overlap Upset plot shows that a very small fraction of these connections are actually shared.

Unlike the previous steps of the pipeline, where were we evaluated the performance of methods on mock datasets, there is no equivalent dataset that contain a set of known interactions for the evaluation of the network inference algorithms. Therefore, we propose the construction of a consensus network (Figure 5C) involving MAGMA, mLDM, SpiecEasi and SparCC. This consensus network is built by merging the p-values generated from bootstraps of the original taxonomy composition table using the Browns p-value combining method [44] (see Methods section). Based on this approach, MiCoNE reports as default output the consensus network, annotated with weights (correlations for SparCC and direct associations for the other methods) for all four methods.

### The default pipeline

The systematic analyses performed in the previous sections clearly show that the choice of tools and parameters can have a big impact on the final co-occurrence network. For some of these choices (e.g. DADA2 vs. deblur) there is no clear metric to establish a best protocol. For other choices, the mock communities provide an opportunity to select combination of parameters that yield more accurate and robust results. Despite this partial degree of assessment, we wish to suggest a combination of tools and parameters that produce networks that are derived from the combination of tools which performed best on the mock communities, and displayed highest robustness to switching to alternative methods. These tools and parameters are chosen as the defaults for the pipeline and are given in Table 1.

**Table 1:** Default tools and parameters for the pipeline

| Process | Tool | Parameters |
| --- | --- | --- |
| Denoising and Clustering | Dada2/Deblur | default |
| Taxonomy Assignment | NCBI with Blast | RefSeq database |
| OTU Processing | Based on statistical power | Dynamic cutoff |
| Network Inference | Consensus method | - |

The recommended tool for the Denoising and Clustering (DC) step (DADA2 or Deblur) were chosen based on their accuracy in recapitulating the reference sequences in mock communities and synthetic data. The choice of the taxonomy reference database in the Taxonomy Assignment (TA) step is dictated largely by the species expected to be present in the sample as well the database used in similar studies if comparison is a goal. Nevertheless, we suggest NCBI RefSeq along with blast+ as the query tool since the database is updated regularly and has a broad collection of taxonomies. The abundance threshold at the OTU Processing (OP) step is determined automatically based on the number of samples and the required statistical power. Finally, we use the Browns p-value combining method on the networks generated using MAGMA, mLDM, SpiecEasi and SparCC to obtain a final consensus network in the Network Inference (NI) step.

Figure 6A shows the default network compared against networks generated by altering one of the steps of the pipeline from the default. These results indicate that the biggest differences in networks occur when the reference database or the network inference algorithm are changed. Furthermore, the L1 distance of networks generated by altering one of the steps of the pipeline from the default against the default network (Figure 6B) shows that the biggest deviations from the default network occur when the TA and NI steps are changed, reinforcing the same results observed in Figure 2. Figure 7 shows the co-occurrence networks inferred for the hard palate for healthy subjects in a periodontal disease study [45] and the healthy stool microbiome in fecal microbial transplant study [35]. These consensus networks were generated using the default tools and parameters from Table 1.

**Figure 6:**
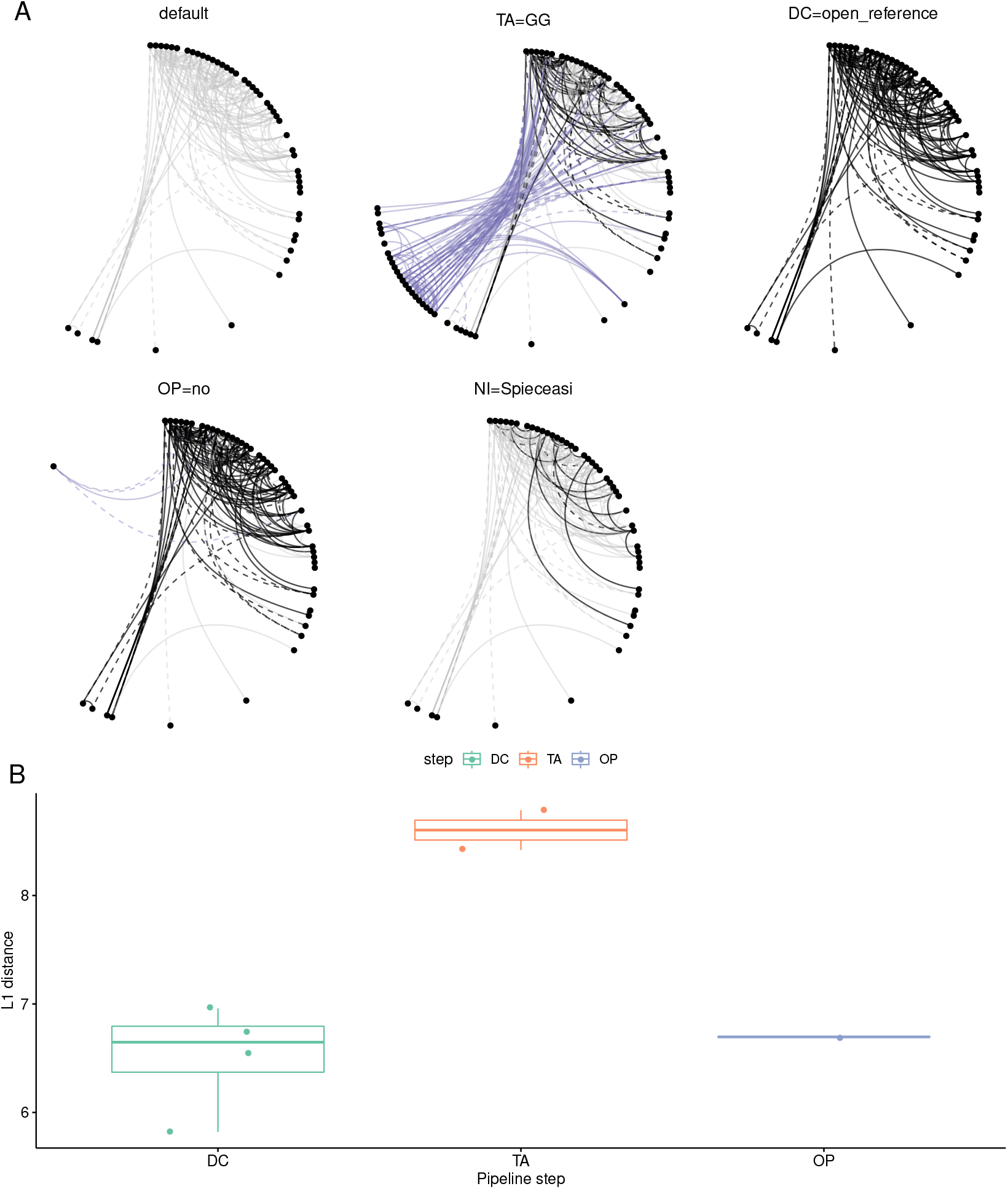
Network inference and taxonomic assignment have the highest influence on the inferred network structures. **(A)** The network constructed using the default pipeline parameters (DC=DADA2, TA=NCBI, OP=on, NI=SparCC) is compared with networks generated when one of the steps use a different tool. The common connections (common with the default network) are in black, connections unique to the network are colored purple and connections in the default network but not present in the current network are gray. **(B)** The L1 distance between the networks generated by changing one step of the default pipeline and the network generated using the default parameters.

**Figure 7:**
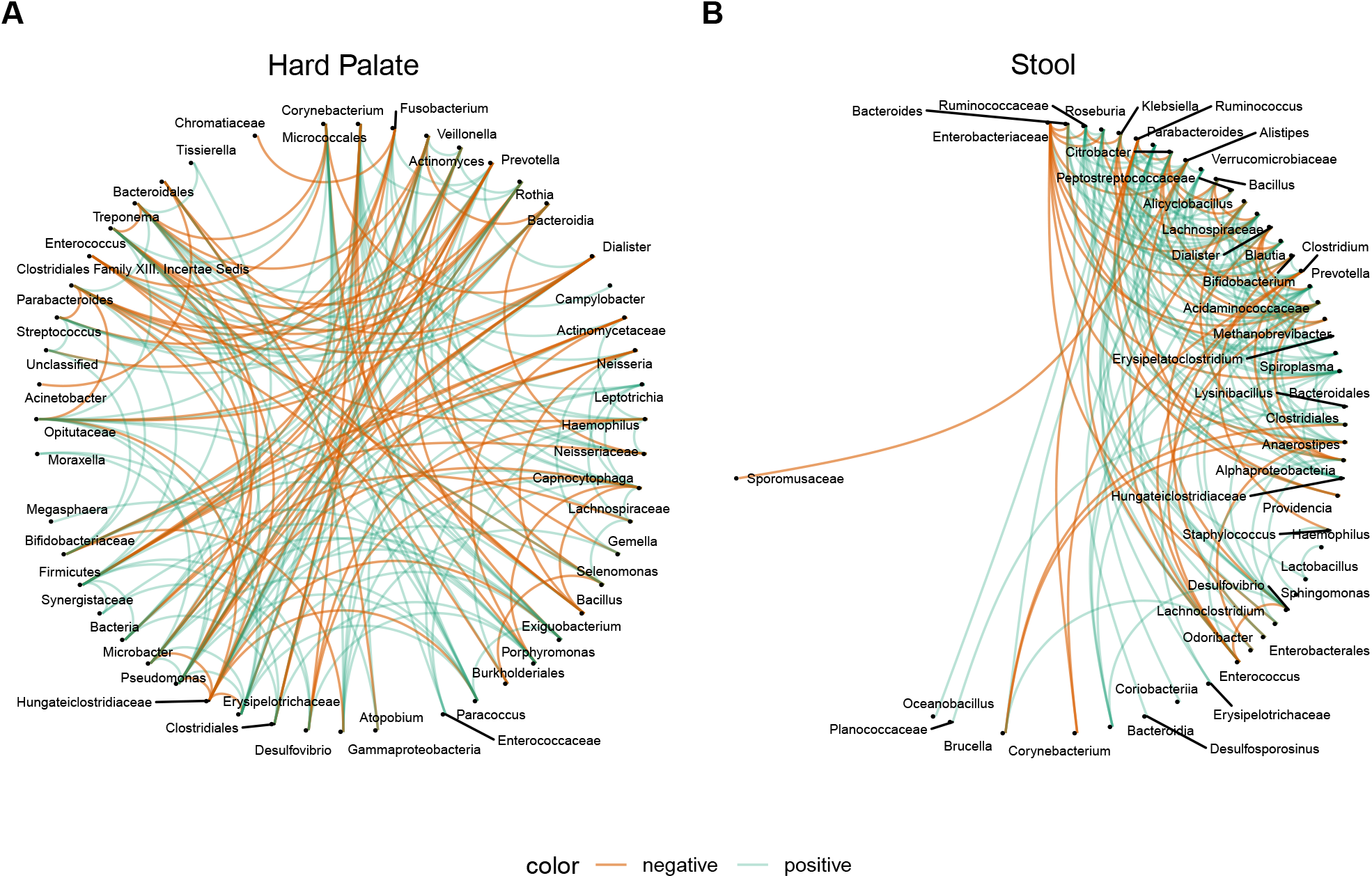
The consensus networks generated using the default pipeline settings. **(A)** Co-occurrence network of the Hard Palate microbiome generated from samples of healthy subjects in a periodontal diseases study. **(B)** Co-occurrence network of the Stool microbiome generated from samples of healthy subjects in a fecal microbiome transplant study.

## Discussion

Co-occurrence associations in microbial communities help identify important interactions that drive microbial community structure and organization. Our analysis shows that networks generated using different combinations of tools and approaches can look significantly different from each other, highlighting the importance of a clear assessment of the source of variability and of tools that provide the most robust and accurate results. Our newly developed integrated software for the inference of co-occurrence networks from 16S rRNA data, MiCoNE, constitutes a freely customizable and user friendly pipeline that allows users to easily test combinations of tools and to compare networks generated by multiple possible choices (see Methods). Importantly, in addition to revisiting the test cases presented in this work, users will be able to explore the effect of various tool combinations on their own datasets of interest. The MiCoNE pipeline is built in a modular fashion. Its plug-and-play architecture will make it possible for users to add new tools and steps, either from existing packages, or from packages that were not examined in the present work, as well as future ones.

The main outcome of this work is thus two-fold: on one hand we transparently reveal the dependence of co-occurrence networks on tool and parameter choices, making it possible to more rigorously assess and compare existing networks. On the other hand, we take advantage of our spectrum of computational options and the availability of mock and synthetic datasets, to suggest a default standard setting, and a consensus approach, likely to yield networks that are robust across multiple tool/parameter choices.

An important caveat related to this last point is the fact that our conclusions are based on the specific datasets used in our analysis. While our datasets cover a relatively broad spectrum of biomes and sequencing pipelines, datasets that have drastically different distributions may require a re-assessment of the best settings through our pipeline.

It is worth pointing out some additional more specific conclusions stemming from the individual steps of our analysis.

The different denoising/clustering methods differ mostly in their identification of sequences that are in low abundances. Hence, they do not have much of an impact on the inferred co-occurrence networks when the sequences of low abundance are removed. However, comparison of inferred and expected reference sequences and their abundances in mock community datasets has allowed us to identify DADA2 as the method which best recapitulates the expected sequence composition. For the current work we have decided to focus on the tools most widely used at the time of the analysis. Some tools that we recently published (e.g. dbOTU3 [46]) as well as older popular methods like mothur [47] have not been included in the study, but could be added into the pipelines in future updated analyses.

The choice of taxonomy database was found to be the most important factor in the inference of a microbial co-occurrence network, contributing ∼ 20% of the total variance. The frequent changes in the taxonomy nomenclature coupled with the frequency of updates to the various 16S reference databases create inherent differences [41] in taxonomy hierarchies in these databases. Our analysis revealed that no particular reference database performs better than the others across all scenarios. We suggest that that choice of the database should be made based on possible reported or inferred biases in the representation of given biomes in a specific databases [41]. The default reference database in the pipeline is the NCBI 16S RefSeq database as it is more frequently updated and is most compatible with the blast+ query tool. We also enable users to use custom databases [48] with the blast+ and naive bayes classifiers that are incorporated into the pipeline (from QIIME2).

Filtering out taxa that are present in low abundances in all samples did not increase (in most datasets tested) the proportion of taxa in common between taxonomy tables generated using different reference databases. However, we do observe that the reduction in the number of taxa leads to better agreement in the networks inferred through different methods. Moreover, filtering is necessary in order to increase the power in tests of significance when the number of taxa is much greater than the number of samples.

The networks generated by different network inference methods show considerable differences in edge-density and connectivity. One reason for this is the underlying assumptions regarding sparsity, distribution and compositionality that the algorithms make. The consensus network created by merging the networks inferred using the different network inference methods enables the creation of a network whose links have evidence based on multiple inference algorithms.

Exploring the effects of these combinations of methods on the resultant networks is difficult and inconvenient since different tools differ in their input and output formats and require inter-converting between the various formats. The pipeline facilitates this comparative exploration by providing a variety of modules for inter-conversion between various formats, and by allowing easy incorporation of new tools as modules.

We envision that MiCoNE, and the underlying tools and databases that help process amplicon sequencing data into co-occurrence networks, will be increasingly useful towards building large comparative analyses across studies. By having a unified transparent tool to compute networks, it will be possible to reprocess available 16S datasets to obtain networks that are directly comparable to each other. Furthermore, even in the analysis of published networks across studies and processing methods, MiCoNE could help understand underlying biases of each network, which could in turn be taken into account upon making cross-study comparisons.

## Materials and Methods

### Datasets

The study uses three kinds of 16S rRNA sequencing datasets: real datasets, mock datasets and synthetic datasets. Real datasets are collections of sequencing reads obtained from naturally occurring microbial community samples. The current study used healthy stool samples from a fecal microbiome transplant study [35] and healthy saliva samples from a periodontal disease study [45] as real datasets for analysis. The mock community 16S datasets are real sequencing data obtained for artificially assembled collections of species in known proportions. The mock datasets used for this study, obtained from mockrobiota [33], are labelled mock4, mock12 and mock16. The mock4 community is composed of 21 bacterial strains. Two replicate samples from mock4 contain all species in equal abundances, and two additional replicate samples contain the same species in unequal abundances. The mock12 community is composed of 27 bacterial strains that include closely related taxa with some pairs having only one to two nucleotide difference from another. The mock16 community is composed of 49 bacteria and 10 archea, all represented in equal amount. The synthetic datasets were generated using an artificial read simulator called ART [34]. Three different microbial composition profiles were used as input; reads were generated using a soil and water microbiome composition profiles from the EMP [2] and healthy gut microbiome project from the fecal microbiome transplant study [35]. The reads are simulated using the NCBI RefSeq database as the reference sequence pool and the “art_illumina” sequence profile with a mutation rate of 2%. The scripts used to generate the synthetic data are in the scripts folder of the repository (https://github.com/segrelab/MiCoNE-pipeline-paper).

### MiCoNE

The flowchart describing the workflow of MiCoNE (Microbial Co-occurrence Network Explorer), our complete 16S data-analysis pipeline, is shown in Figure 1. The pipeline integrates many publicly available tools as well as custom R or Python modules and scripts to extract co-occurrence associations from 16S sequence data. Each of these tools corresponds to a distinct R or python module that recapitulates the relevant analyses. All such individual modules are available as part of the MiCoNE package. The inputs to the pipeline by default are the raw community 16S rRNA sequence reads, but the software can be alternatively configured to use trimmed sequences, OTU tables and other types of intermediate data. The final output of the pipeline is the inferred network of co-occurrence relationships among the microbes present in the samples.

The MiCoNE pipeline provides both a Python API as well as a command-line interface and only requires a single configuration file. The configuration file lists the inputs, output and the steps to be performed during runtime, along with the parameters to be used (if different from defaults) for the various steps. Since the entire pipeline run-through is stored in the form of a text file (the configuration file), subsequent runs are highly reproducible and changes can be easily tracked using version control. It uses the nextflow workflow manager [49] under the hood, making it readily usable on local machines, cluster or cloud with minimal configuration change. It also allows for automatic parallelization of all possible processes, both within and across samples. The pipeline is designed to be modular: each tool or method is organized into modules which can be easily modified or replaced. This modular architecture simplifies the process of adding new tools (refer to modules section in the MiCoNE documentation). The main components of the pipeline are detailed in the subsequent sections.

### Denoising and Clustering (DC)

This module deals with processing the raw 16S sequence data into OTU or ESV count tables. It consists of the following processes: quality control, denoising (or clustering) and chimera checking. The quality control process handles the demultiplexing and quality control steps such as trimming adapters and trimming low-quality nucleotide stretches from the sequences. The denoise/cluster process handles the conversion of the demultiplexed, trimmed sequences into OTU or ESV count tables (some methods, like closed reference and open reference clustering, perform clustering and taxonomy assignment in the same step). The chimera checking process handles the removal of chimeric sequences created during the Polymerase Chain Reaction (PCR) step. The output of this module is a matrix of counts, that describes the number of reads of a particular OTU or ESV (rows of the matrix) present in each sample (columns of the matrix). The options currently available in the pipeline for denoising and clustering are: open reference clustering, closed reference clustering and de novo clustering methods from QIIME1 v1.9.1 [22] and denoising methods from DADA2 v1.14 [23] and Deblur v1.1.0 [36]. The quality filtering and chimera checking tools are derived from those used in QIIME2 v2019.10.0 and DADA2.

### Taxonomy Assignment (TA)

This module deals with assigning taxonomies to either the representative sequences of the OTUs or directly to the ESVs. In order to assign taxonomies to a particular sequence we need a taxonomy database and a query tool. The taxonomy database contains the collection of 16S sequences of micro-organisms of interest and the query tool allows one to compare a sequence of interest to all the sequences in the database to identify the best matches. Finally, a consensus method is used to identify the most probable match from the list of best matches. The pipeline incorporates GG 13_8 [24], SILVA 132 [25] and the NCBI (16S RefSeq as of Oct 2019) [40] databases for taxonomy assignment and the Naive Bayes classifier from QIIME2 and NCBI blast as the query tools (from QIIME2). The consensus algorithm used is the default method used by the classifiers in QIIME2.

### OTU and ESV Processing (OP)

This module deals with normalization, filtering and applying transformations to the OTU or ESV counts matrix. Rarefaction is a normalization technique used to overcome the bias that might arise due to variable sampling depth in different samples. This is performed either by sub-sampling or by normalization of the matrix to the lowest sampling depth [26]. Rarefaction is usually followed by filtering, which is performed to remove samples or features (OTUs or ESVs) from the count matrix that are sparse. In order to determine the filtering threshold we fix the number of samples and correlation detection power needed and determine the number of features to be used. Finally, transformations are performed in order to correct for and overcome the compositional bias that is inherent in a counts matrix (in most cases this is handled by the network inference algorithm).

### Network Inference (NI)

This module deals with the inference of co-occurrence associations from the OTU or ESV counts matrix. These associations can be represented as a network, with nodes representing taxonomies of the micro-organisms and edges representing the association between them. A null model is created by re-sampling and bootstrapping the correlation/interaction matrix and is used to calculate the significance of the inferred associations by calculating the p-values against this null model [50]. The pipeline includes Pearson, Spearman and FastSpar v0.0.10 (a faster implementation of SparCC) [50] as the pairwise correlation metrics, and SpiecEasi v1.0.7 [28], mLDM v1.1 [42] and MAGMA [27] as the direct association metrics. The empirical Browns method [44] is used for combining p-values from the various methods to obtain a consensus p-value, which is used to create the consensus network.

### Network Variability

In order to compare across different networks, and analyze the degree of variability induced by the choice of different modules and parameters, we organized multiple networks into a single mathematical structure that we could use for linear regression. In particular, we transformed the adjacency matrix of each co-occurrence network into a vector. We then merged the networks generated from all possible combinations of tools into a table (N, see below) in which each column represents one network.

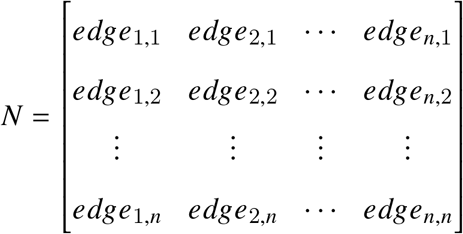

In other words, *N* is the merged table, each column *N*_*i*_ is the vector representation of one of the networks, and each row *L*_*i*_ represents the one particular edge in all networks (assigned 0 if the edge does not exist in the network).

We use linear regression to express each link *L*_*i*_ as a linear function of categorical variables that describe the possible options in each of the first three steps of the pipeline.

In particular, we infer parameters *α*_*i*_ such that:

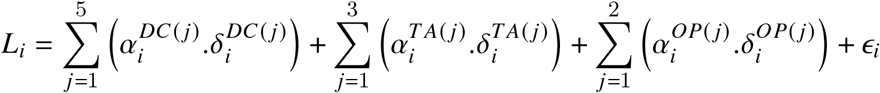

where, *α*_*i*_ are the coefficients of the regression, *ϵ*_*i*_ are the residuals and *δ*_*i*_ are the indicator variables that correspond to the processes utilized in the pipeline used to create the network *N*_*i*_; for example, 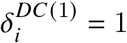 if the DC(1) process was used in the generation of the network *N*_*i*_. Here, (i) DC(1) = “closed reference”, DC(2) = “open reference”, DC(3) = “de novo”, DC(4) = “dada2”, DC(5) = “deblur” (ii) TA(1) = “GreenGenes”, TA(2) = “SILVA”, TA(3) = “NCBI” (iii) OP(1) = “no filtering”, OP(2) = “filtering”.

The variance contributed by each step of the pipeline is calculated for every connection in the merged table through ANOVA using the Python statsmodels package and is shown in Figure 2B. The total variance for the network is calculated by adding the variances for each connection. The PCA analysis is also performed on the merged table to generate Figure 2C.

## Code and Data Availability

Pipeline: https://github.com/segrelab/MiCoNE

Data and scripts: https://github.com/segrelab/MiCoNE-pipeline-paper

## Acknowledgments

We are grateful to members of the Segrè lab for helpful discussions and for feedback on the manuscript. This work was partially funded by grants from the National Institutes of Health (National Institute of General Medical Sciences, award R01GM121950; National Institute of Dental and Craniofacial Research, award number R01DE024468; and National Institute on Aging, award number UH2AG064704), the U.S. Department of Energy, Office of Science, Office of Biological & Environmental Research through the Microbial Community Analysis and Functional Evaluation in Soils SFA Program (m-CAFEs) under contract number DE-AC02-05CH11231 to Lawrence Berkeley National Laboratory, the National Science Foundation (grants 1457695 and NSFOCE-BSF 1635070), the Human Frontiers Science Program (RGP0020/2016), and the Boston University Interdisciplinary Biomedical Research Office. KSK was supported by Simons Foundation Grant #409704, by the Research Corporation for Science Advancement through Cottrell Scholar Award #24010, by the Scialog grant #26119, and by the Gordon and Betty Moore Foundation grant #6790.08.

## Contributions

Designed the research project: DK, KK, DS, ZH, CDL. Performed analysis: DK, GB. Wrote the first draft of the manuscript: DK. Revised and wrote final version of the manuscript: DK, DS, KK.

## Supplementary

**Table S1:**
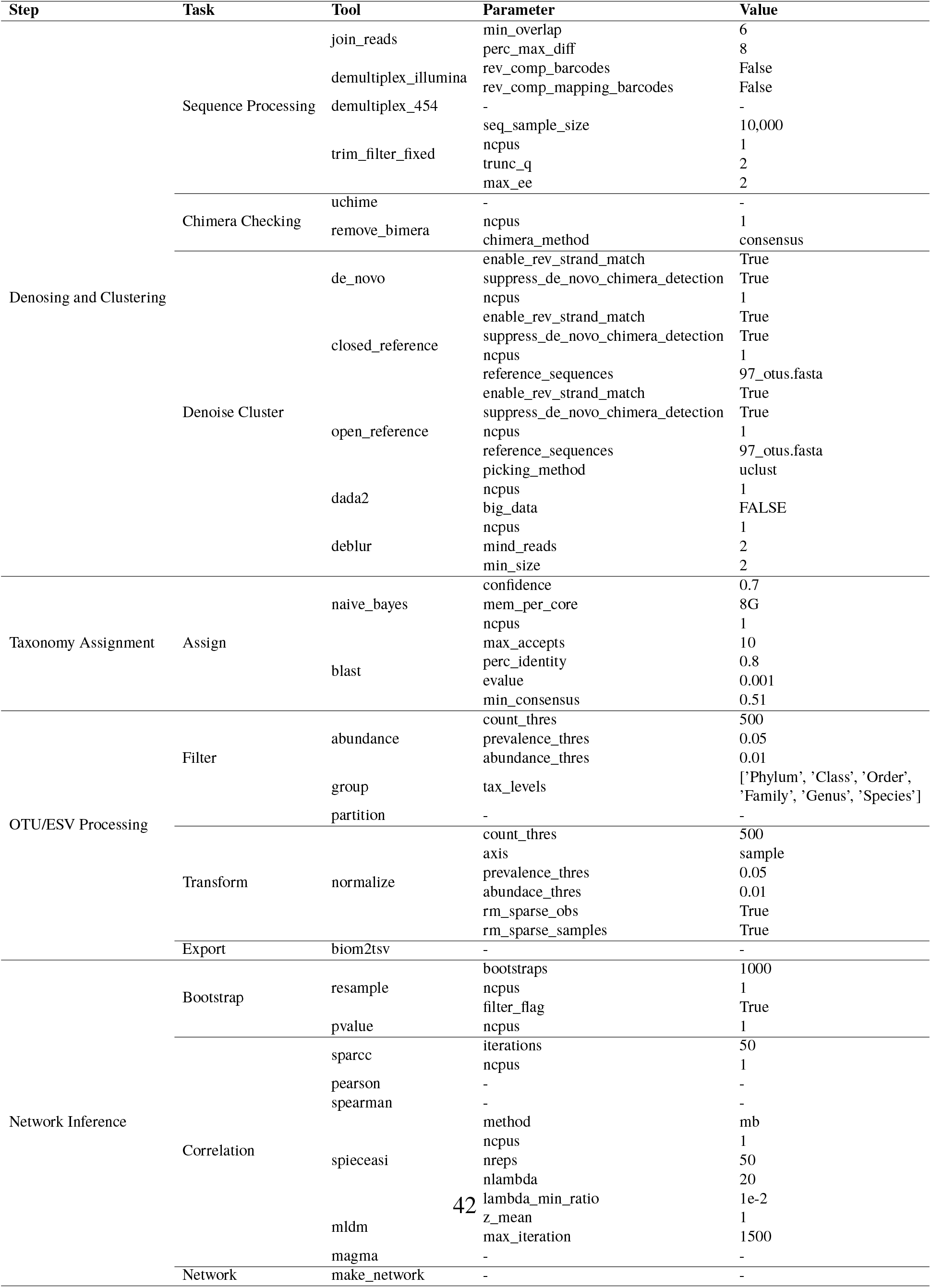
The default parameters used in the various tools of the pipeline

---

**Figure S1:**
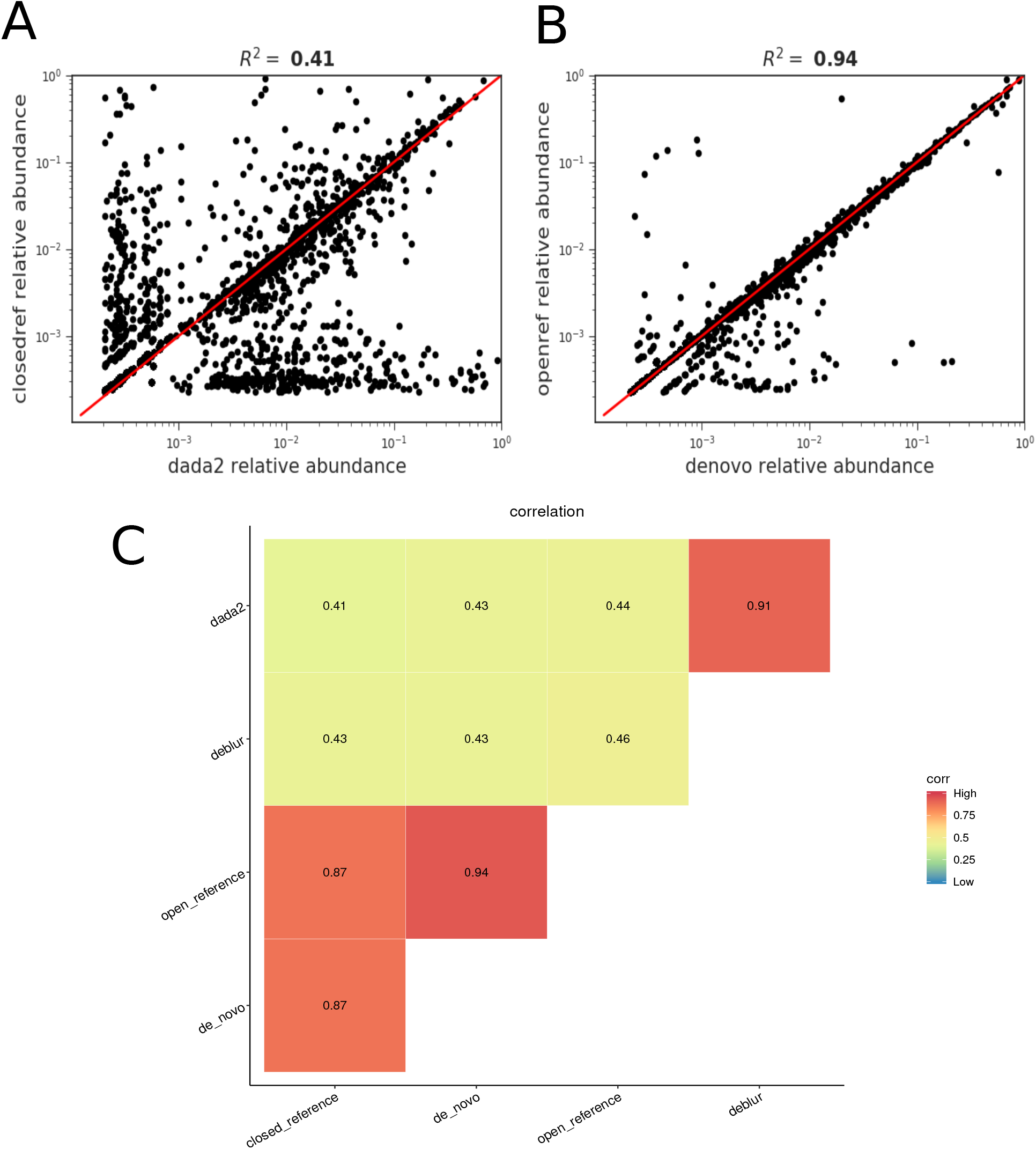
Comparison of various denoising and clustering algorithms used in the pipeline. (A, B) Correlation of the abundances of the taxa that are in common between the count matrices created by two different methods. (A) The worst correlation (least similar methods) is between open-reference and dada2. (B) The best correlation (most similar methods) is between open-reference and denovo. (C) A heatmap showing the R^2^ of all pairwise comparisons of the methods.

**Figure S2:**
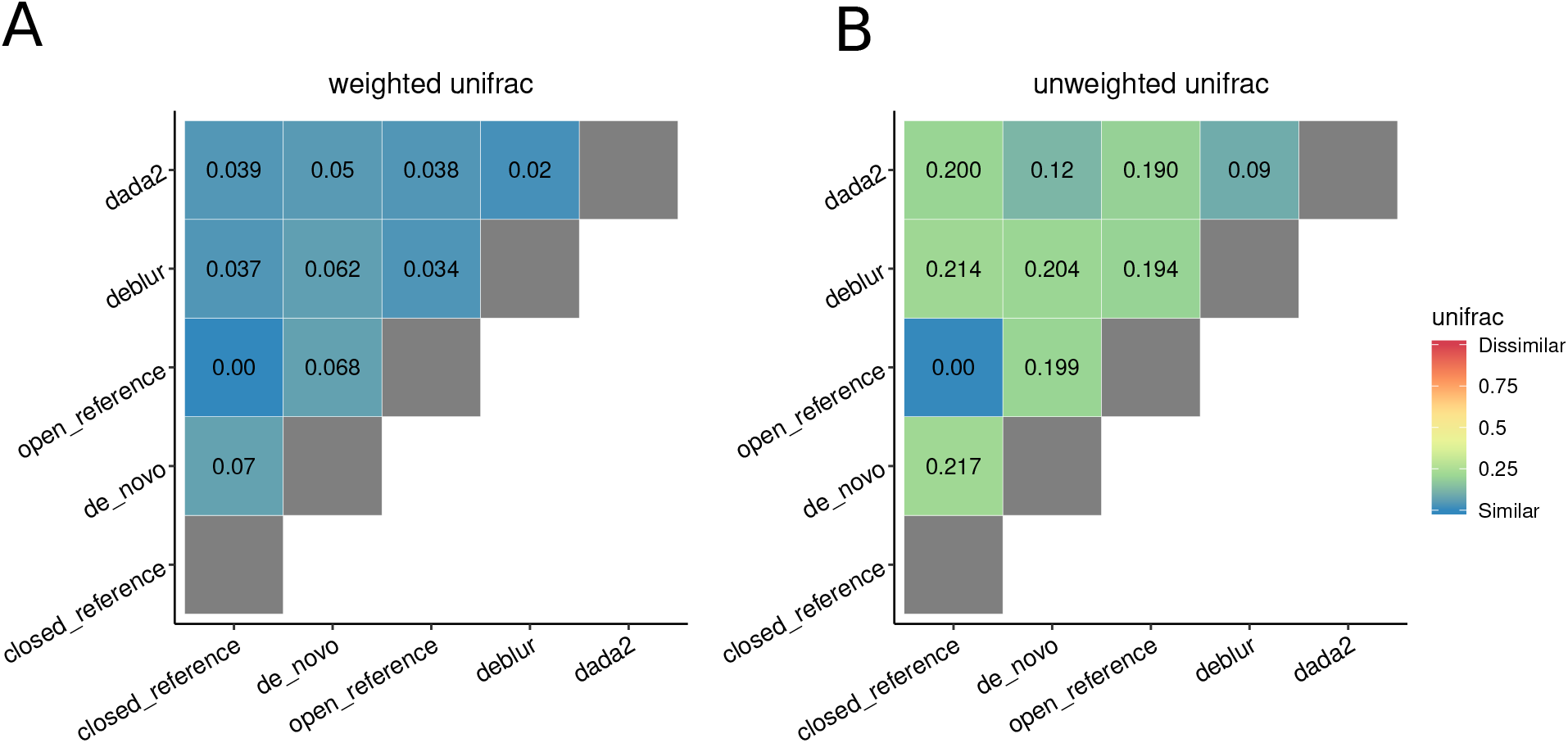
Heatmaps showing the weighted and unweighted unifrac distances for the hard palate dataset analysis. (A) weighted unifrac distances and (B) unweighted unifrac distances between the representative sequences generated by different denoising and clustering algorithms. These results are in agreement with the stool microbiome dataset.

**Figure S3:**
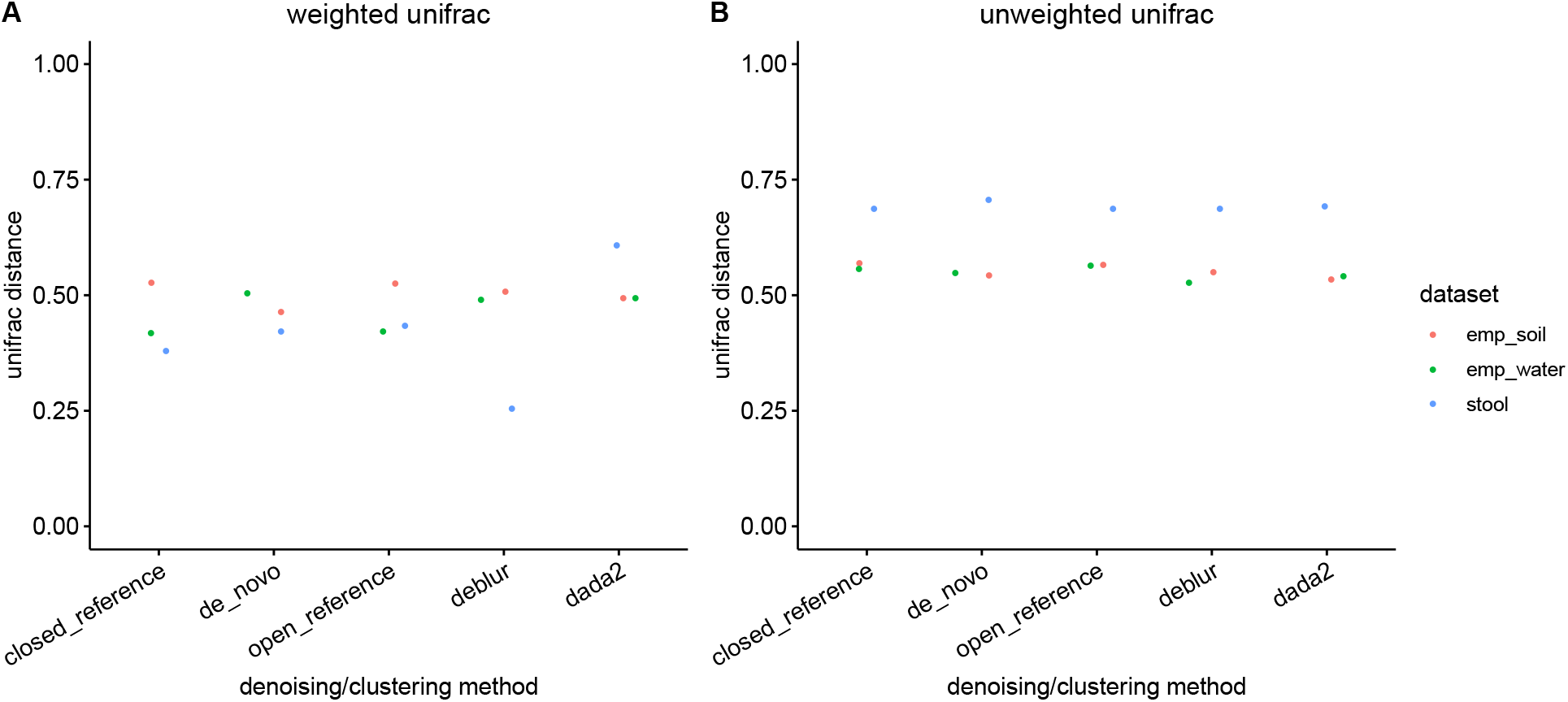
The distributions of the average weighted UniFrac distance between the expected sequence profile and the calculated sequence profile in the synthetic datasets. We observe no significant difference between the various methods on the synthetic datasets used for this study.

**Figure S4:**
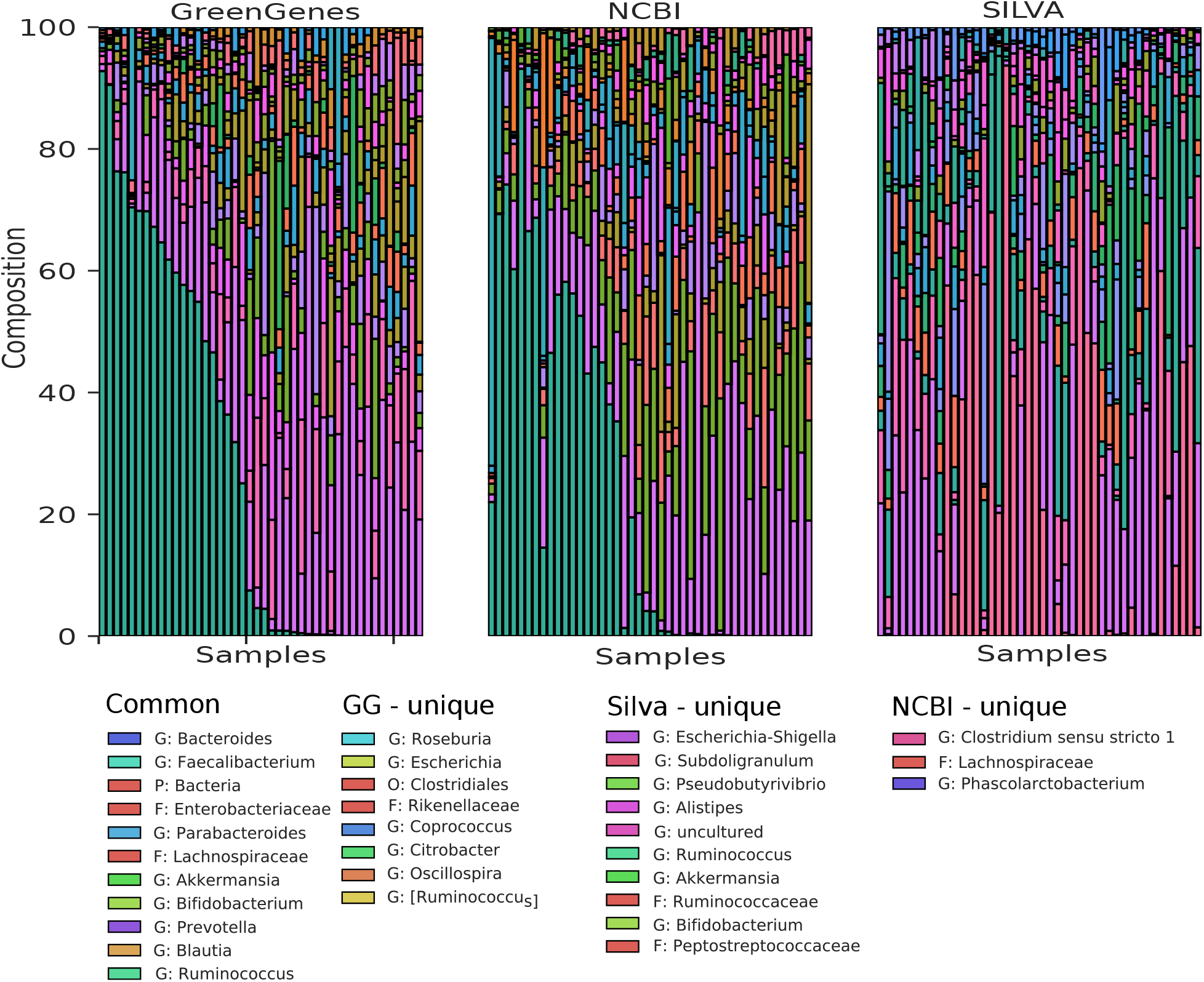
**(A)** Taxonomy composition of the 20 most abundant genera predicted for the stool microbiome dataset generated using different taxonomy references databases: Greengenes, SILVA and NCBI. The legend shows the common and the unique genera among the taxonomy assignments.

**Figure S5:**
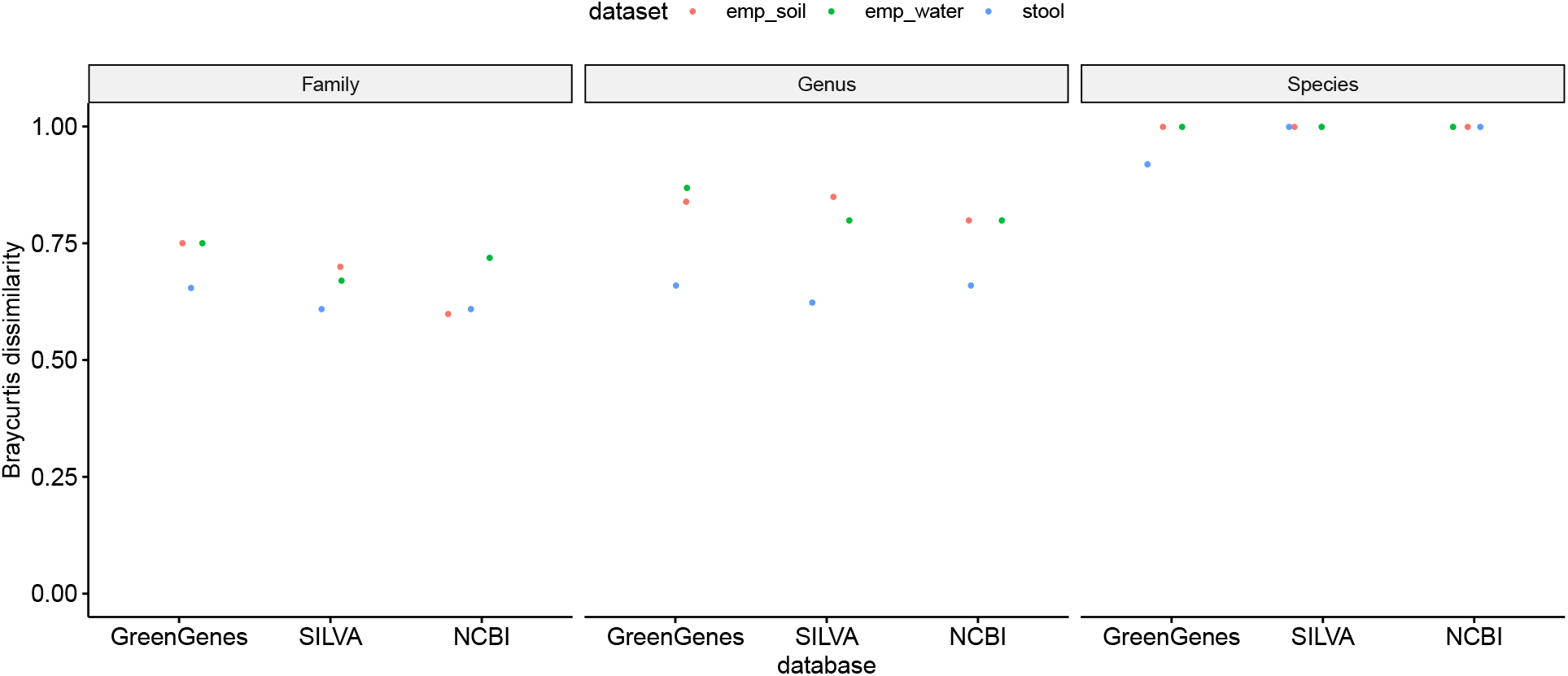
The bray-curtis dissmilarity between the expected taxonomic composition and generated taxonomic composiion for the synthetic datasets.

**Figure S6:**
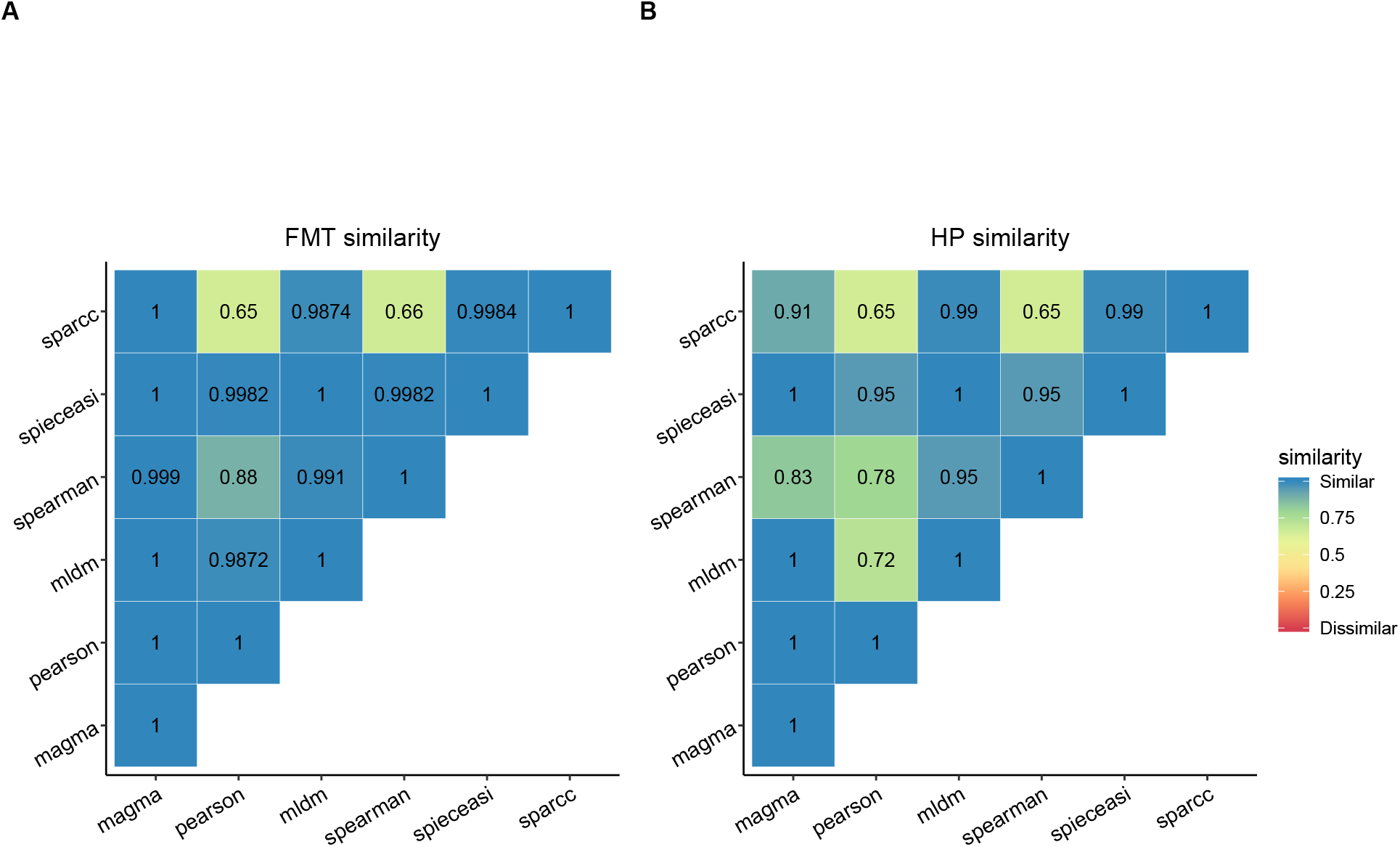
The similarity between the networks generated using the different network inference algorithms for stool dataset (A) and the hard palate dataset (B). The similarity between the various methods was found to vary with the dataset used.

**Figure S7:**
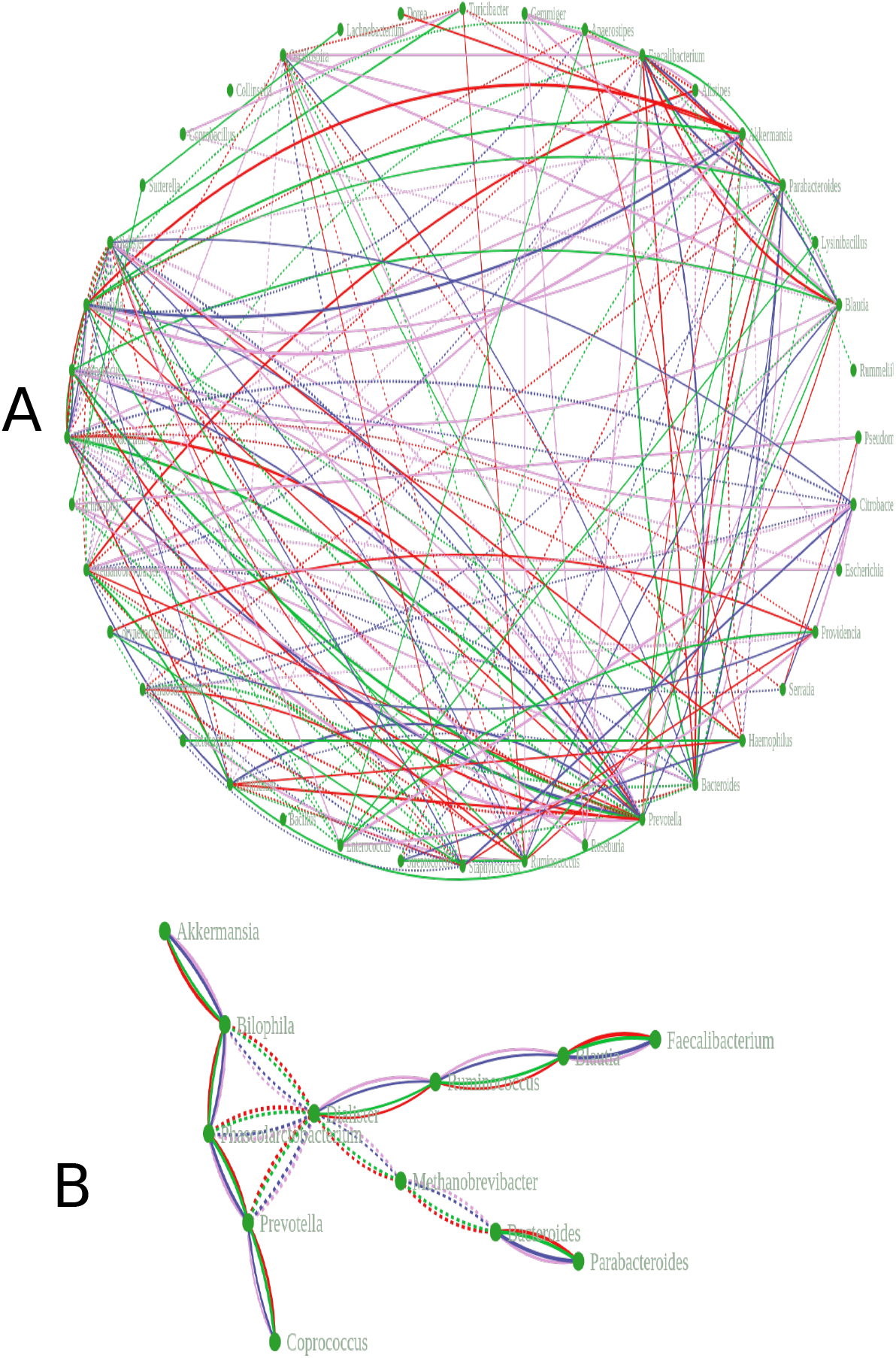
A network showing union (A) and intersection (B) of networks generated using different denoising and clustering tools on the Stool dataset.

## References

[1] Melanie Ghoul and Sara Mitri. “The Ecology and Evolution of Microbial Competition.” In: Trends in microbiology 24.10 (Oct. 2016), pp. 833–845. ISSN: 1878-4380. DOI: 10.1016/j.tim.2016.06.011. URL: http://www.ncbi.nlm.nih.gov/pubmed/27546832.

[2] Luke R. Thompson et al. “A communal catalogue reveals Earth’s multiscale microbial diversity”. In: Nature 551.7681 (Nov. 2017), p. 457. ISSN: 0028-0836. DOI: 10.1038/nature24621. URL:http://www.nature.com/doifinder/10.1038/nature24621.

[3] Takashi Narihiro and Yoichi Kamagata. “Genomics and Metagenomics in Microbial Ecology: Recent Advances and Challenges.” In: Microbes and environments 32.1 (2017), pp. 1–4. ISSN: 1347-4405. DOI: 10.1264/jsme2.ME3201rh. URL: http://www.ncbi.nlm.nih.gov/pubmed/28367917%20http://www.pubmedcentral.nih.gov/articlerender.fcgi?artid=PMC5371069.

[4] Juan Jovel et al. “Characterization of the Gut Microbiome Using 16S or Shotgun Metagenomics.” In: Frontiers in microbiology 7 (2016), p. 459. ISSN: 1664-302X. DOI: 10.3389/fmicb.2016.00459. URL: http://www.ncbi.nlm.nih.gov/pubmed/27148170%20http://www.pubmedcentral.nih.gov/articlerender.fcgi?artid=PMC4837688.

[5] Jason Lloyd-Price, Galeb Abu-Ali, and Curtis Huttenhower. “The healthy human microbiome.” In: Genome medicine 8.1 (2016), p. 51. ISSN: 1756-994X. DOI: 10.1186/s13073-016-0307-y. URL: http://www.ncbi.nlm.nih.gov/pubmed/27122046%20http://www.pubmedcentral.nih.gov/articlerender.fcgi?artid=PMC4848870.

[6] Houjin Zhang and Kang Ning. “The Tara Oceans Project: New Opportunities and Greater Challenges Ahead.” In: Genomics, proteomics & bioinformatics 13.5 (Oct. 2015), pp. 275–7. ISSN: 2210-3244. DOI: 10.1016/j.gpb.2015.08.003. URL: http://www.ncbi.nlm.nih.gov/pubmed/26546828%20http://www.pubmedcentral.nih.gov/articlerender.fcgi?artid=PMC4678785.

[7] Barbara A. Human Microbiome Project Consortium et al. “A framework for human microbiome research.” In: Nature 486.7402 (June 2012), pp. 215–21. ISSN: 1476-4687. DOI: 10.1038/nature11209. URL: http://www.ncbi.nlm.nih.gov/pubmed/22699610%20http://www.pubmedcentral.nih.gov/articlerender.fcgi?artid=PMC3377744.

[8] Rajita Menon, Vivek Ramanan, and Kirill S. Korolev. “Interactions between species introduce spurious associations in microbiome studies”. In: PLOS Computational Biology 14.1 (Jan. 2018). Ed. by Stefano Allesina, e1005939. ISSN: 1553-7358. DOI: 10.1371/journal.pcbi.1005939. URL: http://dx.plos.org/10.1371/journal.pcbi.1005939.

[9] Lisa Röttjers and Karoline Faust. “From hairballs to hypotheses–biological insights from microbial networks”. In: FEMS Microbiology Reviews 42.6 (Nov. 2018), pp. 761–780. ISSN: 1574-6976. DOI: 10.1093/femsre/fuy030. URL: https://academic.oup.com/femsre/article/42/6/761/5061627.

[10] Jack A. Gilbert et al. “Microbiome-wide association studies link dynamic microbial consortia to disease”. In: Nature 535.7610 (2016), pp. 94–103. ISSN: 14764687. DOI: 10.1038/nature18850. arXiv: NIHMS150003.

[11] Baohong Wang et al. “The Human Microbiota in Health and Disease”. In: Engineering 3.1 (Feb. 2017), pp. 71–82. ISSN: 20958099. DOI: 10.1016/J.ENG.2017.01.008. URL: http://linkinghub.elsevier.com/retrieve/pii/S2095809917301492.

[12] José E Belizário and Mauro Napolitano. “Human microbiomes and their roles in dysbiosis, common diseases, and novel therapeutic approaches.” In: Frontiers in microbiology 6 (2015), p. 1050. ISSN: 1664-302X. DOI: 10.3389/fmicb.2015.01050. URL: http://www.ncbi.nlm.nih.gov/pubmed/26500616%20http://www.pubmedcentral.nih.gov/articlerender.fcgi?artid=PMC4594012.

[13] Noah Fierer. Embracing the unknown: Disentangling the complexities of the soil microbiome. Oct. 2017. DOI: 10.1038/nrmicro.2017.87. URL: https://pubmed.ncbi.nlm.nih.gov/28824177/.

[14] Shuo Jiao, Weimin Chen, and Gehong Wei. “Resilience and assemblage of soil microbiome in response to chemical contamination combined with plant growth”. In: Applied and Environmental Microbiology 85.6 (Mar. 2019). ISSN: 10985336. DOI: 10.1128/AEM.02523-18. URL: http://aem.asm.org/.

[15] Ryan H. Hsu et al. “Microbial Interaction Network Inference in Microfluidic Droplets”. In: Cell Systems 9.3 (Sept. 2019), 229–242.e4. ISSN: 24054720. DOI: 10.1016/j.cels.2019.06.008. URL: http://www.cell.com/article/S2405471219302315/fulltext%20http:s//www.cell.com/article/S2405471219302315/abstract%20https://www.cell.com/cell-systems/abstract/S2405-4712(19)30231-5.

[16] Xingjin Jian et al. “Microbial microdroplet culture system (MMC): An integrated platform for automated, high-throughput microbial cultivation and adaptive evolution”. In: Biotechnology and Bioengineering 117.6 (June 2020), pp. 1724–1737. ISSN: 0006-3592. DOI: 10.1002/bit.27327. URL: https://onlinelibrary.wiley.com/doi/abs/10.1002/bit.27327.

[17] Steven A Wilbert, Jessica L Mark Welch, and Gary G Borisy. “Spatial Ecology of the Human Tongue Dorsum Microbiome”. In: CellReports 30 (2020), 4003–4015.e3. DOI: 10.1016/j.celrep.2020.02.097. URL: https://doi.org/10.1016/j.celrep.2020.02.097.

[18] Cristal Zuñiga, Livia Zaramela, and Karsten Zengler. “Elucidation of complexity and prediction of interactions in microbial communities”. In: Microbial Biotechnology 10.6 (2017), pp. 1500–1522. DOI: 10.1111/1751-7915.12855.

[19] Jonathan Friedman and Eric J. Alm. “Inferring Correlation Networks from Genomic Survey Data”. In: PLoS Computational Biology 8.9 (Sept. 2012). Ed. by Christian von Mering, e1002687. ISSN: 1553-7358. DOI: 10.1371/journal.pcbi.1002687. URL: http://dx.plos.org/10.1371/journal.pcbi.1002687.

[20] Richa Bharti and Dominik G Grimm. “Current challenges and best-practice protocols for microbiome analysis”. In: Briefings in Bioinformatics 2019.00 (Dec. 2019), pp. 1–16. ISSN: 1477-4054. DOI: 10.1093/bib/bbz155. URL: https://academic.oup.com/bib/advance-article/doi/10.1093/bib/bbz155/5678919.

[21] Jolinda Pollock et al. The madness of microbiome: Attempting to find consensus “best practice” for 16S microbiome studies. Apr. 2018. DOI: 10.1128/AEM.02627-17. URL: https://doi.org/10.1128/AEM.02627-17.

[22] J Gregory Caporaso et al. “QIIME allows analysis of high-throughput community sequencing data”. In: Nature Methods 7.5 (May 2010), pp. 335–336. ISSN: 1548-7091. DOI: 10.1038/nmeth.f.303. URL: http://www.nature.com/articles/nmeth.f.303.

[23] Benjamin J Callahan et al. “DADA2: High-resolution sample inference from Illumina amplicon data”. In: Nature Methods 13.7 (July 2016), pp. 581–583. ISSN: 1548-7091. DOI: 10.1038/nmeth.3869. URL: http://www.nature.com/articles/nmeth.3869.

[24] T Z DeSantis et al. “Greengenes, a chimera-checked 16S rRNA gene database and workbench compatible with ARB.” In: Applied and environmental microbiology 72.7 (July 2006), pp. 5069–72. ISSN: 0099-2240. DOI: 10.1128/AEM.03006-05. URL: http://www.ncbi.nlm.nih.gov/pubmed/16820507%20http://www.pubmedcentral.nih.gov/articlerender.fcgi?artid=PMC1489311.

[25] Christian Quast et al. “The SILVA ribosomal RNA gene database project: improved data processing and web-based tools”. In: Nucleic Acids Research 41.D1 (Nov. 2012), pp. D590–D596. ISSN: 0305-1048. DOI: 10.1093/nar/gks1219. URL: http://academic.oup.com/nar/article/41/D1/D590/1069277/The-SILVA-ribosomal-RNA-gene-databaseproject.

[26] Sophie J Weiss et al. “Effects of library size variance, sparsity, and compositionality on the analysis of microbiome data”. In: PeerJ PrePrints 3 (2015), e1408. ISSN: 2167-9843. DOI: 10.7287/peerj.preprints.1157v1. arXiv: peerj.preprints.270v1 [10.7287]. URL: https://doi.org/10.7287/peerj.preprints.1157v1%7B%5C%%7D5Cnhttps://peerj.com/preprints/1157v1/%7B%5C#%7Dsupp-8.

[27] Arnaud Cougoul, Xavier Bailly, and Ernst C Wit. “MAGMA: inference of sparse microbial association networks”. In: (2019). DOI: 10.1101/538579. URL: https://doi.org/10.1101/538579.

[28] Zachary D. Kurtz et al. “Sparse and Compositionally Robust Inference of Microbial Ecological Networks”. In: PLOS Computational Biology 11.5 (May 2015). Ed. by Christian von Mering, e1004226. ISSN: 1553-7358. DOI: 10.1371/journal.pcbi.1004226. URL: http://dx.plos.org/10.1371/journal.pcbi.1004226.

[29] Kevin P. Keegan, Elizabeth M. Glass, and Folker Meyer. “MG-RAST, a Metagenomics Service for Analysis of Microbial Community Structure and Function”. In: Humana Press, New York, NY, 2016, pp. 207–233. DOI: 10.1007/978-1-4939-3369-3_13. URL: http://link.springer.com/10.1007/978-1-4939-3369-3%7B%5C_%7D13.

[30] Qiita-open-source microbial study management platform. URL: https://qiita.ucsd.edu/(visited on 05/22/2018).

[31] Jonathan L. Golob et al. “Evaluating the accuracy of amplicon-based microbiome computational pipelines on simulated human gut microbial communities”. In: BMC Bioinformatics 18.1 (2017), p. 283. ISSN: 1471-2105. DOI: 10.1186/s12859-017-1690-0. URL: http://bmcbioinformatics.biomedcentral.com/articles/10.1186/s12859-017-1690-0.

[32] Sophie Weiss et al. “Correlation detection strategies in microbial data sets vary widely in sensitivity and precision”. In: Isme J 10.7 (2016), pp. 1–13. ISSN: 1751-7362. DOI: 10.1038/ismej.2015.235. URL: http://dx.doi.org/10.1038/ismej.2015.235%7B%5C%%7D5Cn10.1038/ismej.2015.235.

[33] Nicholas A Bokulich et al. “mockrobiota: a Public Resource for Microbiome Bioinformatics Benchmarking.” In: mSystems 1.5 (2016). ISSN: 2379-5077. DOI: 10.1128/mSystems.00062-16. URL: http://www.ncbi.nlm.nih.gov/pubmed/27822553%20http://www.pubmedcentral.nih.gov/articlerender.fcgi?artid=PMC5080401.

[34] Weichun Huang et al. “ART: a next-generation sequencing read simulator.” In: Bioin-xsformatics (Oxford, England) 28.4 (Feb. 2012), pp. 593–4. ISSN: 1367-4811. DOI: 10.1093/bioinformatics/btr708. URL: http://www.ncbi.nlm.nih.gov/pubmed/22199392%20http://www.pubmedcentral.nih.gov/articlerender.fcgi?artid=PMC3278762.

[35] Dae-Wook Kang et al. “Microbiota Transfer Therapy alters gut ecosystem and improves gastrointestinal and autism symptoms: an open-label study”. In: Microbiome 5.1 (Dec. 2017), p. 10. ISSN: 2049-2618. DOI: 10.1186/s40168-016-0225-7. URL: http://microbiomejournal.biomedcentral.com/articles/10.1186/s40168-016-0225-7.

[36] Amnon Amir et al. “Deblur Rapidly Resolves Single-Nucleotide Community Sequence Patterns.” In: mSystems 2.2 (Apr. 2017), e00191–16. ISSN: 2379-5077. DOI: 10.1128/mSystems.0019-16. URL: http://www.ncbi.nlm.nih.gov/pubmed/28289731%20http://www.pubmedcentral.nih.gov/articlerender.fcgi?artid=PMC5340863.

[37] Catherine A Lozupone et al. “Quantitative and qualitative beta diversity measures lead to different insights into factors that structure microbial communities.” In: Applied and environmental microbiology 73.5 (Mar. 2007), pp. 1576–85. ISSN: 0099-2240. DOI: 10.1128/AEM.01996-06. URL: http://www.ncbi.nlm.nih.gov/pubmed/17220268%20http://www.pubmedcentral.nih.gov/articlerender.fcgi?artid=PMC1828774.

[38] Catherine Lozupone and Rob Knight. “UniFrac: a new phylogenetic method for comparing microbial communities.” In: Applied and environmental microbiology 71.12 (Dec. 2005), pp. 8228–35. ISSN: 0099-2240. DOI: 10.1128/AEM.71.12.8228-8235.2005. URL: http://www.ncbi.nlm.nih.gov/pubmed/16332807%20http://www.pubmedcentral.nih.gov/articlerender.fcgi?artid=PMC1317376.

[39] Jacob T. Nearing et al. “Denoising the Denoisers: an independent evaluation of microbiome sequence error-correction approaches”. In: PeerJ 6 (Aug. 2018), e5364. ISSN: 2167-8359. DOI: 10.7717/peerj.5364. URL: https://peerj.com/articles/5364.

[40] Eric W Sayers et al. “Database resources of the National Center for Biotechnology Information.” In: Nucleic acids research 37.Database issue (Jan. 2009), pp. D5–15. ISSN: 1362-4962. DOI: 10.1093/nar/gkn741. URL: http://www.ncbi.nlm.nih.gov/pubmed/18940862%20http://www.pubmedcentral.nih.gov/articlerender.fcgi?artid=PMC2686545.

[41] Monika Balvočiūtė and Daniel H. Huson. “SILVA, RDP, Greengenes, NCBI and OTT — how do these taxonomies compare?” In: BMC Genomics 18.S2 (Mar. 2017), p. 114. ISSN: 1471-2164. DOI: 10.1186/s12864-017-3501-4. URL: http://bmcgenomics.biomedcentral.com/articles/10.1186/s12864-017-3501-4.

[42] Yuqing Yang, Ning Chen, and Ting Chen. “Inference of Environmental Factor-Microbe and Microbe-Microbe Associations from Metagenomic Data Using a Hierarchical Bayesian Statistical Model.” In: Cell systems 4.1 (Jan. 2017), 129–137.e5. ISSN: 2405-4712. DOI: 10.1016/j.cels.2016.12.012. URL: http://www.ncbi.nlm.nih.gov/pubmed/28125788.

[43] Alexander Lex et al. UpSet: Visualization of Intersecting Sets. Tech. rep. URL: http://grouplens.org/datasets/movielens/.

[44] William Poole et al. “Combining dependent P-values with an empirical adaptation of Brown’s method”. In: (). DOI: 10.1093/bioinformatics/btw438. URL: https://www.bioconduc.

[45] Casey Chen et al. “Oral microbiota of periodontal health and disease and their changes after nonsurgical periodontal therapy”. In: The ISME Journal 12.5 (May 2018), pp. 1210–1224. ISSN: 1751-7362. DOI: 10.1038/s41396-017-0037-1. URL: http://www.nature.com/articles/s41396-017-0037-1.

[46] Scott W. Olesen, Claire Duvallet, and Eric J. Alm. “DbOTU3: A new implementation of distribution-based OTU calling”. In: PLoS ONE 12.5 (2017), pp. 1–13. ISSN: 19326203. DOI: 10.1371/journal.pone.0176335.

[47] Patrick D Schloss et al. “Introducing mothur: open-source, platform-independent, community-supported software for describing and comparing microbial communities.” In: Applied and environmental microbiology 75.23 (Dec. 2009), pp. 7537–41. ISSN: 1098-5336. DOI: 10.1128/AEM.01541-09. URL: http://www.ncbi.nlm.nih.gov/pubmed/19801464%20http://www.pubmedcentral.nih.gov/articlerender.fcgi?artid=PMC2786419.

[48] Jarmo Ritari et al. In: BMC Genomics 6.1 (Dec. 2011), p. 1056. ISSN: 1471-2164. DOI: 10.1186/s12864-015-2265-y.

[49] Paolo Di Tommaso et al. “Nextflow: A tool for deploying reproducible computational pipelines”. In: F1000Research 4 (July 2015). DOI: 10.7490/F1000RESEARCH.1110183.1. URL: https://f1000research.com/posters/4-430.

[50] Stephen C Watts et al. “FastSpar: rapid and scalable correlation estimation for compositional data”. In: Bioinformatics (Aug. 2018). Ed. by Oliver Stegle. ISSN: 1367-4803. DOI: 10.1093/bioinformatics/bty734. URL: https://academic.oup.com/bioinformatics/advance-article/doi/10.1093/bioinformatics/bty734/5086389.

